# Mutual homeostasis of charged proteins

**DOI:** 10.1101/2023.08.21.554177

**Authors:** Rupert Faraway, Neve Costello Heaven, Holly Digby, Oscar G. Wilkins, Anob M. Chakrabarti, Ira A. Iosub, Lea Knez, Stefan L. Ameres, Clemens Plaschka, Jernej Ule

## Abstract

Protein dosage is regulated to maintain cellular homeostasis and health. The dosage of proteins containing disordered low complexity domains (LCDs) must be particularly well-controlled to prevent aberrant disease, yet no mechanism to maintain homeostasis has been identified^1, 2^. Here we report a mutual homeostatic mechanism that controls the concentration of such proteins, termed ’interstasis’, in which proteins with similar LCDs co-regulate their combined dosage through collective negative feedback. We focused on the mechanism that exploits the fundamental multivalency of GA-rich RNA regions that encode charged LCDs, including those with arginine-enriched mixed charge domains (R-MCDs). Modest variations in the abundance of an R-MCD protein change the properties of nuclear speckles, a protein-RNA condensate, selectively trapping multivalent GA-rich mRNAs to promote their nuclear retention. This interstasis depends on conserved codon biases, shared by amniotes, which enhance the multivalency of GA-rich regions encoding charged LCDs. The threshold of interstasis is modulated by CLK kinases, which affect the nuclear speckle localisation of proteins such as TRA2B, key binder of GA-rich RNAs. Notably, many classes of LCDs are encoded by RNA regions containing multivalency-enhancing codon biases, each preferentially bound by specific proteins, suggesting that interstasis might co-regulate many classes of functionally related LCD-containing proteins through dose-sensitivity of various types of protein-RNA condensates.

## Main

Low complexity domains (LCDs) of proteins, which contain overrepresentations of specific amino acids, are exceptionally common in proteins across all domains of life^3^. LCD-containing proteins have an increased propensity to localise into biomolecular condensates, the formation of which is highly sensitive to the concentration of these proteins^4^. It has been demonstrated through fitness screens in multiple eukaryotic species that proteins with disordered LCDs are more dosage sensitive^1^. These proteins often display an increased propensity for aggregation that is a hallmark of many diseases, especially in neurodegeneration^4–6^. It is therefore important to understand how the dosage of LCD-containing proteins is regulated and homeostatically controlled.

The properties of many types of condensate are often influenced by groups of proteins containing similar LCDs; for instance, arginine-rich mixed charge domains (R-MCDs) contribute to the condensation of nuclear speckles and acidic LCDs to the nucleolus^7, 8^. This selectivity is a result of high incidence of residues in LCDs that promote disorder and that form weak protein-protein or protein-RNA interactions which contribute to the specificity of each of these condensates for a class of LCD-containing proteins^5, 9–11^. This suggests that the homeostasis of large classes of LCDs with similar properties may need to be mutually co-regulated.

Proteins with disordered LCDs have been observed to show below-average protein abundance and above-average RNA abundance, indicating that their dosage regulation may occur post-transcriptionally and post-translationally^1, 2^. Known mechanisms exist for feedback regulation of global protein or RNA concentrations^12, 13^ and for auto-regulation of individual transcription factors or RBPs^14, 15^. However, it has remained unclear if and how LCD-containing proteins can be more selectively regulated as functionally related groups in response to changes in their dosage.

Soon after the discovery of the genetic code, it was observed that amino acids with similar characteristics tend to have similar codons^16^ which has been proposed to mitigate the effects of genetic mutations or translation errors^17, 18^. As LCDs are defined as over-representations of amino acids, often with similar characteristics, regions of coding sequences (CDS) that encode LCDs are thus expected to contain repetitions of similar codons. These regions of RNA should therefore contain abundant similar RNA sequence motifs, often referred to as multivalent RNA regions.

In the present study, we find that LCDs are encoded by multivalent regions of RNA, which we find bound by specific assemblies of RNA-binding proteins (RBPs). Notably, LCD-encoding CDS regions are not multivalent solely due to the characteristics of the genetic code, but also due to strong, conserved codon biases. To demonstrate the importance of this property, we focus on the multivalent GA-rich mRNAs that encode charged LCD-containing proteins. We show that a dose-sensitive response of nuclear speckles to the concentration of R-MCDs, a sub-type of charged LCD, leads to selective retention of this multivalent mRNA in the nuclear speckle, preventing their export and translation. Therefore, an RNA-dependent mechanism drives the mutual homeostasis of the dosage of charged LCD-containing proteins, a phenomenon which we term interstasis.

### Multivalent RNA regions encode LCDs

Previously developed tools to identify multivalent RNA motifs bound by RNA-binding proteins have focused on analysis of specific experimental data, such as to identify sets of motifs enriched in specific co-regulated RNAs, or bound by specific RBPs^19–21^. To enable the analysis of multivalency potential in RNA sequences not constrained to a specific RBP or motif set, we designed a scoring algorithm that produces a generalised RNA multivalency (GeRM) score for each k-mer in any RNA sequence (Figure 1A).

**Figure 1:**
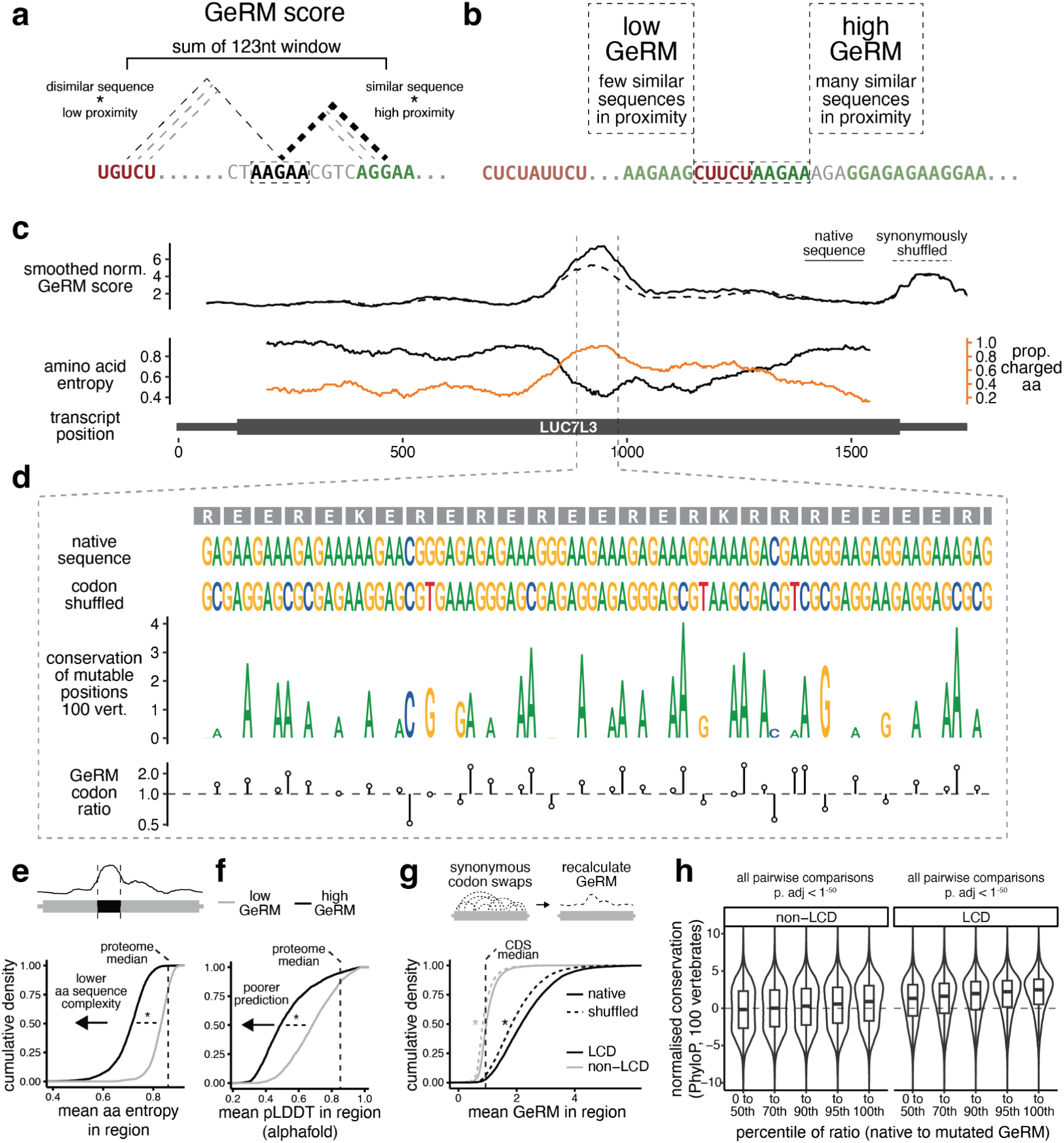
Generalised RNA multivalency scoring identifies codon biased multivalent regions that encode low complexity domains. a, A schematic describing the calculation of a GeRM score for an individual 5-mer. b, An example of two 5-mers with either high or low GeRM scores. c, An example of a transcript from the gene LUC7L3. The smoothed GeRM score is shown along the transcript in the upper panel (solid line), with the dashed line depicting the average smoothed GeRM score after synonymous codon shuffling. The amino acid entropy of the encoded sequence is shown in the lower panel (black line), while the proportion of charged amino acids in that window is shown in the orange line. d, The native DNA and amino acid sequences of LUC7L3 within the GeRM peak. A codon shuffled sequence in which all codons must be synonymously shifted is shown. The conservation across 100 vertebrates (PhyloP) for each position that can tolerate synonymous mutation is shown, with the height of the letter corresponding to the PhyloP score. Below, a ratio is shown that compares the GeRM scores associated with the native codon choice to the GeRM scores associated with any possible synonymous mutation. e, The mean entropy for amino acid sequences encoded by high GeRM CDS regions (black line) or encoded outside of high GeRM CDS regions, but within the same protein (grey line). The median amino acid entropy across the entire proteome is shown by the dashed line. f, As in panel e, but comparing the AlphaFold-predicted pLDDT values inside (black line) and outside (grey line) protein regions encoded by high GeRM CDS regions. g, The mean GeRM scores within CDS regions encoding LCDs (black lines), or the rest of the CDS (grey lines). The mean GeRM scores in those regions after synonymously reordering the codons within each transcript (dashed lines). h, The normalised conservation across 100 vertebrates of synonymously mutable positions in coding sequences that either encode LCDs or do not. Codons are binned by the degree that the native codon choice supports sequence multivalency, where codons with the highest ratio support the multivalency the most. Conservation values are normalised to the middle position of the codon, which is never synonymously mutable, and each codon is normalised to have a median conservation value of 0. All pairwise statistical comparisons performed using a Welch t-test with FDR correction for multiple testing, where * = p < 1^-15^.

For a given k-mer at each position along a transcript, the GeRM score is calculated by taking all k-mers in a defined window around the given k-mer. The similarity of these surrounding k-mers to the given k-mer is calculated by taking the inverse exponent of their Hamming distance. This similarity is weighted based on the proximity to the given k-mer, and the weighted similarity scores are then summed for each k-mer (Figure 1A). Thus, k-mers that have a large number of similar k-mers nearby receive a high GeRM score, indicating high multivalency potential (Figure 1B). Focusing on mRNA sequences, we applied the GeRM algorithm to the longest coding transcript for every human gene. For example, the GeRM score profile of LUC7L3 mRNA reveals a region with high GeRM score in the middle of the coding sequence (CDS) where large numbers of similar GA-rich k-mers are found in close proximity (Figure 1C, D).

We reasoned that coding regions with high RNA multivalency would likely encode stereotyped amino acids and therefore LCDs. In the case of LUC7L3, the high GeRM region encodes exclusively charged amino acids (Figure 1C, D). To assess this, we first systematically identified the regions of high GeRM scores in the CDS (“GeRM regions”) by identifying the regions where the smoothed GeRM scores passed the 98th percentile threshold. We then calculated the information entropy of amino acid sequences in a sliding window and found that the entropy of sequences encoded by GeRM regions was significantly lower compared to the remaining sequences of the same proteins, as well as to the average throughout the proteome (Figure 1E). Moreover, we used AlphaFold2 prediction confidence to approximate protein disorder and found that GeRM regions preferentially encode regions of proteins that are predicted to be disordered (Figure 1F). These observations suggest that multivalent coding sequences preferentially encode disordered LCDs.

### Codon bias in LCDs promotes RNA multivalency

One reason for the multivalency of LCD-encoding RNA regions could be the characteristics of the genetic code, as amino acids with similar characteristics have similar codons^16^. In the example of LUC7L3, codon choices reinforce the A-richness of the region (Figure 1C-D), while in the example of CCDC61, codon choices reinforce the G-richness (Figure S1A-B). We asked if biased usage of codons in these regions might further promote their multivalency potential. To test this, we synonymously shuffled the codons within each transcript and calculated the mean GeRM potential across ten shuffles (Figure 1C, G). We then defined LCDs as regions in the bottom 2% of amino acid entropy. The GeRM potential in regions encoding LCDs was markedly higher than in the rest of the CDS, but synonymously shuffling the order of the codons reduced the GeRM potential significantly (Figure 1G). This suggests that, especially in regions encoding LCDs, codon choices tend to promote the multivalency potential of the RNA.

We speculated that if codon usage supports high multivalency of sequences encoding LCDs, then these codon usage biases would tend to be evolutionarily conserved. To test this, we tested every possible synonymous codon substitution in the transcriptome and calculated the change in GeRM potential of each overlapping 5-mer (Figure S1C). Next, we compared the GeRM potentials of the native k-mers and the mutated k-mers. When the ratio of native to mutated GeRM potential is high, then the native codon usage promotes the local multivalency potential. The average ratio for all but four possible synonymously mutable codons was positive, with more common codons having higher ratios on average (Figure S1D). To account for the general differences in amino acid composition and conservation of LCDs (Figure S1F), we normalised the conservation of each synonymously mutable codon and then compared the conservation of the synonymously mutable nucleotide (typically the wobble position) to the conservation of the middle nucleotide in each codon, which can never be synonymously mutated. We found that the more strongly a native codon promotes multivalency, the more conserved the codon is across mammals or vertebrates, and that this effect is especially strong within the regions that encode LCDs (Figure 1H, S1E, S1G). This analysis suggests that evolutionary selection pressure tends to preserve codons that enhance high multivalency potential of RNA sequences encoding LCDs.

### Charged domains are encoded by GA-multivalent RNA regions

To understand in more detail how the sequence characteristics of GeRM regions are linked to those of the encoded LCDs, we calculated the contribution that each 5-mer made to the total GeRM score of each region, reduced the dimensionality of the data with UMAP and clustered the regions located within CDS using HDBSCAN. We observed diverse clusters, some of which represented trinucleotide repeats, and others without such stereotyped repetition: a GC-rich cluster, a C-rich cluster, and three GA-rich clusters, one with more adenines, one with more guanines, and one with interspersed cytosines (Figure 2A, S2A). To understand how these multivalent classes are linked to various types of LCDs that they might encode, we determined the most common amino acids encoded within each CDS GeRM region. We found that GeRM regions with similar multivalent RNA motifs tend to encode LCDs that are dominated by similar amino acids (Figure 2B, S2B). We confirmed that all types of GeRM region encoded low complexity amino acid sequences and that their native codon usage supported their multivalency (Figure S2D, S2E). Interestingly, the GA-rich GeRM regions encoded domains rich in charged amino acids, with the ratio of adenine to guanine in the multivalent motifs biasing the encoded domain towards positive or negative charge, respectively.

**Figure 2:**
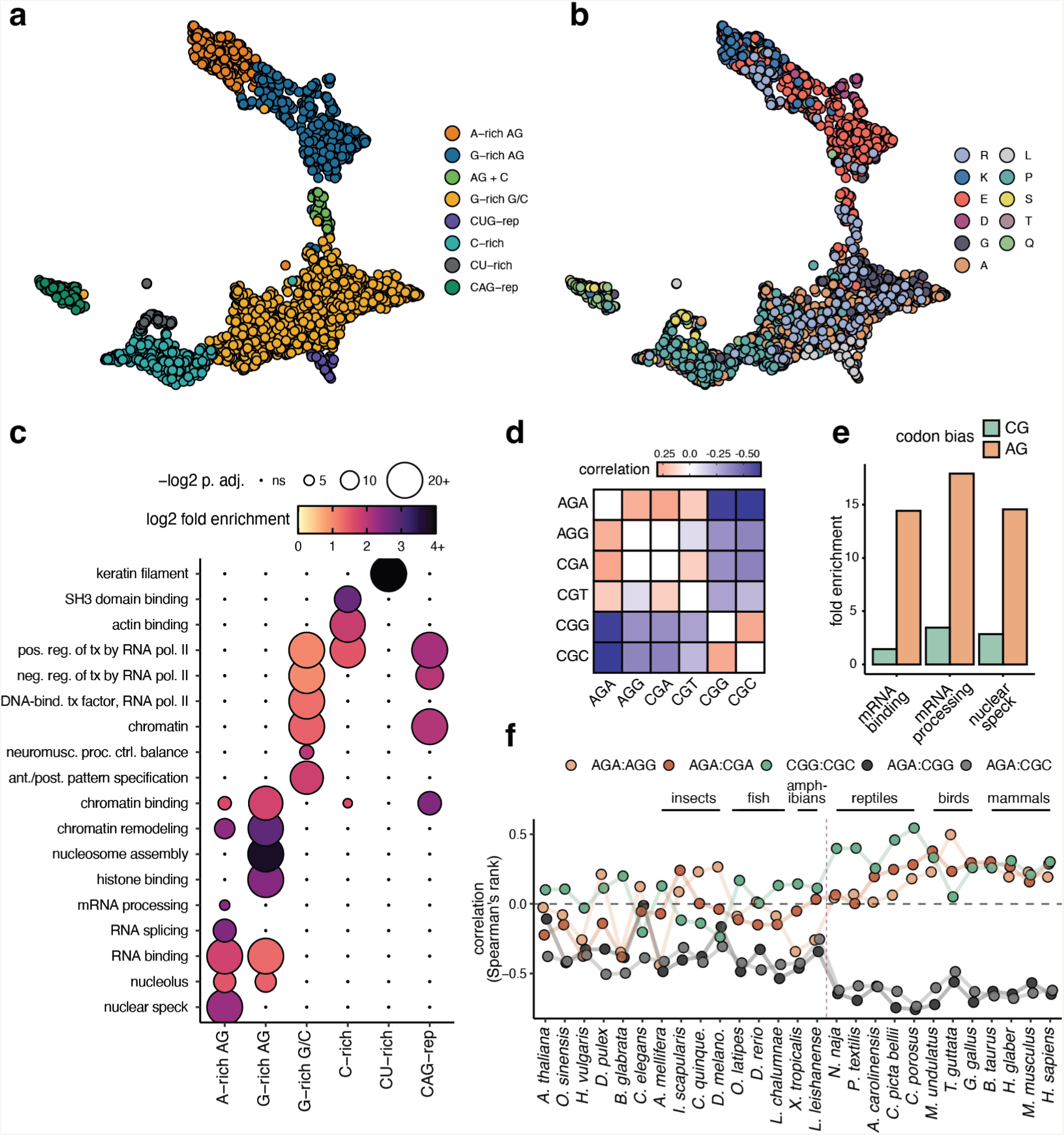
GeRM subtypes encode LCDs with distinct properties and functions. a, A UMAP embedding in which each point is a high GeRM CDS region and is assigned to a cluster. b, UMAP embedding from 2a, with each GeRM region recolored to show the most common amino acid encoded within the region. c, A heatmap of gene ontology fold-enrichments (colour) and significance values (size) for genes containing GeRM regions from the clusters shown in 2a. d, The correlation (Spearman’s rank) of arginine codon usage within LCDs containing at least 20% arginine (R-LCDs). e, The gene ontology fold-enrichments for genes containing R-LCDs encoded by GA-rich codons or CG rich codons. f, Analysis of codon usage correlations within R-LCDs in different species. Species to the right of the dashed vertical line are amniotes.

Given the relationship between the types of GeRM regions and the encoded LCDs, we asked to what extent the GeRM region types are enriched in specific protein functions. We performed an ontology analysis of gene sets containing each type of GeRM region and found that each set was associated with specific terms (Figure 2C). The GA-rich GeRM regions were primarily associated with gene expression-related terms, with GC-rich and CAG-repeat GeRM regions enriched for transcription-related terms, with both types of GA-rich regions enriched for RNA-binding and nucleolar terms, and with G-rich GA-rich regions additionally enriched for chromatin-related terms. The C-rich GeRM regions were most enriched in SH3 domain and actin-binding terms, whereas the CU-rich regions were enriched for keratin filament genes. Therefore, each type of multivalent CDS feature is enriched in functionally related groups of transcripts.

Arginine-rich domains were common in various types of LCDs encoded by GeRM regions, primarily GA-rich or GC-rich regions (Fig 2B, S2C). Arginine can be encoded by six different codons. Therefore, it is not surprising that its codons can be found within diverse multivalent RNA sequences. What is unknown, however, is whether various types of arginine codons are biased for different types of LCDs. To assess such LCD-linked bias, we calculated the arginine codon usage within each LCD that contained at least 20% arginine (referred to as R-LCDs). We observed a positive correlation between AGA, AGG and CGA arginine codons, and between CGG and CGC codons, and strong anti-correlations between these two groups, indicating that the two classes of codons tend to co-occur in different classes of R-LCDs (Figure 2D). We found a strong principal component separating R-LCDs into those where arginines were encoded primarily by GA-rich or CG-rich codons, and we classified these R-LCDs based on this codon-usage principal component (Figure S3A, S3B).

When we compared the gene ontology enrichments of R-LCD genes with either GA- or CG-rich arginine codons, we found that GA-biased R-LCDs were much more highly enriched for nuclear speckle, mRNA processing, and RNA binding terms (Figure 2E). The R-LCDs with GA-rich arginine codons contained significantly more charged residues of either positive or negative charge (Figure S3C, S3F), while those with CG-rich codons contained more proline, alanine and glycine (Figure S3C), suggesting that GA-rich arginine codons are preferred in R-MCDs. We previously found that GA-rich multivalent regions preferentially encode charged amino acids while CG-rich multivalent regions encode proline, alanine and glycine residues. Therefore, arginine codon choice within GeRM regions conforms to the multivalent identity of the region as a whole, such that functionally distinct classes of R-LCD are encoded by RNA regions with distinct multivalent identities.

We then asked at what point in evolution this codon bias in R-LCDs emerged. We analysed R-LCDs in a diverse range of species and observed the correlations and anti-correlations between AG and CG arginine codons. While the total proportions of R-LCDs did not dramatically change across species (Figure S3G-H), we observed that the correlations between CG-rich codons, between GA-rich codons, and the anti-correlation between these groups was stable from humans to reptiles, but that this pattern was much less pronounced in amphibians and other species that diverged earlier in evolutionary history (Figure 2F, S3I). In vertebrates, the separation of R-LCDs by codon bias is most pronounced in amniotes, marking the transition to terrestrial animals.

### The interstasis of charged proteins

Previous work demonstrated a role of R-MCDs in speckle condensation, finding that overexpression of an artificial, positively-charged R-MCD sequence resulted in larger nuclear speckles with slower recovery from photobleaching, as well as a greater sequestration of SR proteins into the speckle^8^. This change in the biophysical properties of the speckle was accompanied by a dose-dependent retention of poly-(A)+ mRNA, with the highest expressing cells displaying a near complete retention of poly-(A)+ mRNA in the nuclear speckle. Thus, large increases in the total abundance of R-MCD proteins could be deleterious by preventing nuclear export of all newly-made mRNAs, which underscores the necessity for their homeostatic control.

We investigated whether R-MCD proteins are under homeostatic control, and if the GA-rich GeRM regions, a subset of which encode R-MCDs, play a role in such homeostasis. Homeostatic self-regulation would take place if the collective concentration of R-MCDs could selectively control the export of mRNAs containing GA-rich GeRM regions. In such a case, increased concentration of R-MCD proteins would selectively retain GA-rich mRNAs encoding R-MCD proteins until the homeostasis is re-established. Alternatively, a homeostatic loop could exist in which the concentration of R-MCD proteins antagonises the definition of GA-rich exons.

To address these possibilities, we designed a reporter system in which an R-MCD fused to mScarlet is encoded with variable codon choices and variable intron number. To achieve this, we assembled a pool of genes encoding a fusion of mScarlet and the C-terminal of Peptidylprolyl Isomerase G (PPIG), which contains a characteristic R-MCD encoded by a GA-rich GeRM region (PPIG_402-684,_henceforth PPIG_LCD_). The R-MCD sequence was generated by Gibson assembly of eight variable fragments, such that the same protein sequence was encoded by a library of reporter genes that differed in their degree of GA-rich multivalency, and in the number of constitutive introns (Figure 3A). This was achieved by two alternative choices for each fragment: GA-rich vs. GA-poor codon choice, and intron-containing vs. intronless choice, producing four variations for each fragment (Figure 3A). Since previous work has implicated transcript GC content in the steady state nuclear-cytoplasmic abundances, we ensured that the GC content of the CDS was kept constant ^22^. Due to the constraints on GC content and splicing efficiency, the most highly multivalent sequence was marginally less multivalent than the original RNA sequence that encodes PPIG_LCD_, whereas the least multivalent sequence was more multivalent than most random CDSs of the same length (Figure S4A). Gibson assembly yielded a pool of approximately 45,000 uniquely barcoded plasmids as identified by targeted sequencing (Figure S4C), with a good representation of all possible fragments (Figure S4B). We transfected this plasmid pool into HeLa cells, isolated the RNA, and confirmed with long-read sequencing that all the introns in the reporter were efficiently spliced (Figure S4D).

**Figure 3:**
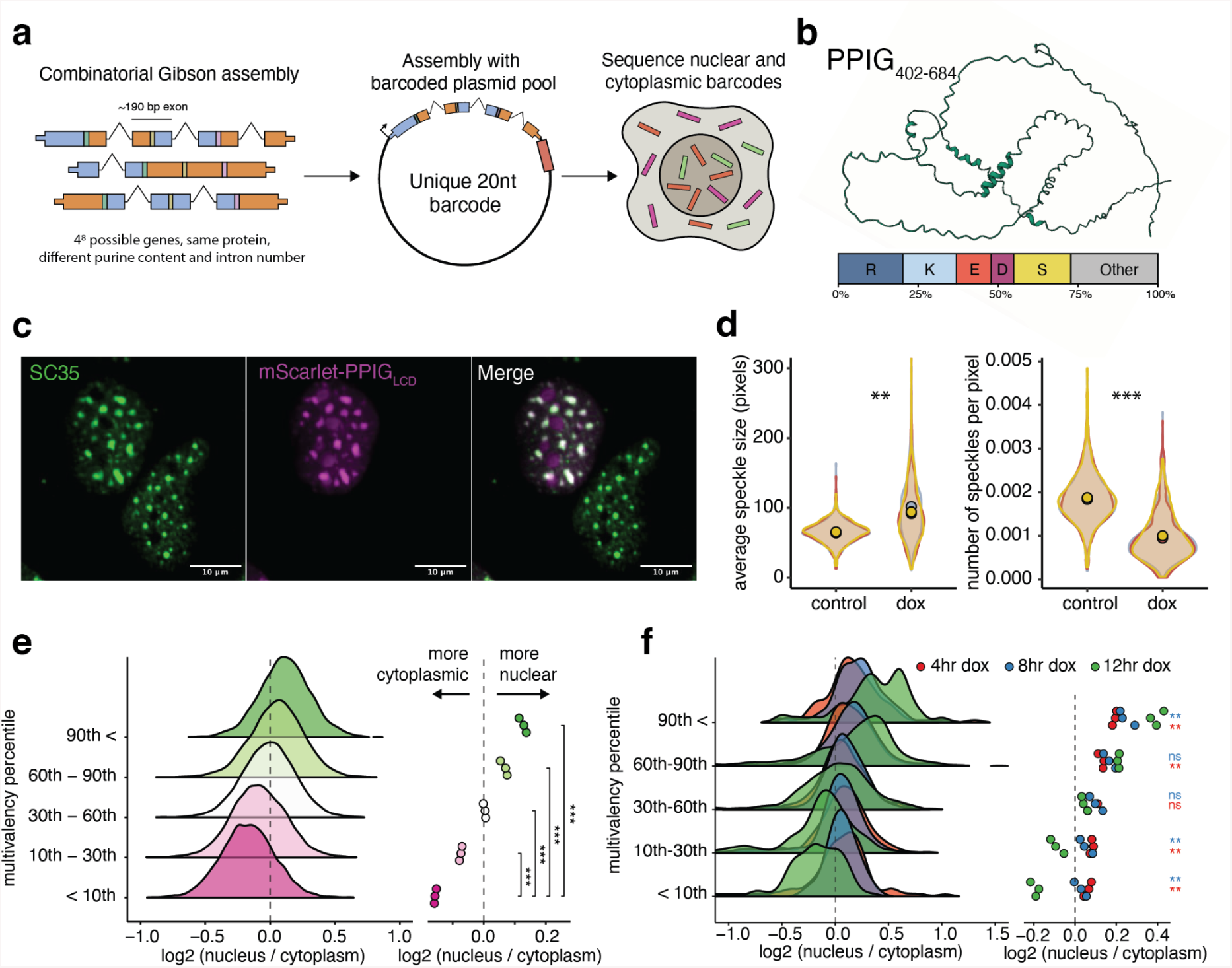
A combinatorially assembled multivalency reporter assessing R-MCD interstasis. a, Schematic describing the assembly of the reporter library. b, AlphaFold prediction of the PPIG_LCD_ structure and the representation of amino acids within the domain. c, Co-localisation of mScarlet-PPIG_LCD_ with nuclear speckles. d, Changes in the size and number of speckles per nucleus after expression of mScarlet-PPIG_LCD_ across 3 replicates, with at least 100 nuclei per replicate. Scale bars represent 10µm. e, The nuclear-cytoplasmic distribution of reporter transcripts binned by their total GeRM scores. Data from cells transfected with the reporter plasmid pool for 16 hours. The right panel shows the group means per sample. f, The nuclear-cytoplasmic abundance ratio of reporter transcripts depending on their sequence multivalency and different lengths of expression induction via doxycycline. The distribution of ratios is shown via ridgeline plot, while the individual group means for each replicate are shown as a dot-plot. All statistical comparisons performed using a Welch t-test with FDR correction for multiple testing, where * = p < 0.05, ** = P < 0.01, *** = P < 0.001.

We cloned the reporter pool into a doxycycline-inducible PiggyBac vector and generated a stable pooled cell line expressing the library of reporter constructs. After 16 hours of doxycycline induction, we observed mScarlet-PPIG_LCD_ localisation to the nucleus of HeLa cells, where it colocalised with SRRM2, a core nuclear speckle component (Figure 3C). Interestingly, expression of PPIG_LCD_ resulted in fewer, larger speckles per nucleus, without reducing the intensity of SRRM2 staining (Figure 3D), consistent with previously reported effects of R-MCDs on speckle condensation properties ^8^. We then transfected HeLa cells with the reporter pool, harvested cells after 16 hours, collected nuclear and cytoplasmic fractions and performed targeted sequencing of the plasmid barcodes present in each fraction. The GeRM score of the reporter construct was strongly correlated with the nuclear-cytoplasmic localisation, such that greater GA-rich multivalency of reporters led to biased mRNA retention in the nuclear fraction (Figure 3E). Interestingly, the number of introns in the reporter transcripts had very little influence on the nuclear-cytoplasmic localisation of the transcripts (Figure S5A). Thus, the reporter mRNAs containing GA-rich GeRM regions were preferentially retained in the nucleus and this retention scaled with the degree of GA-rich multivalency.

If the nuclear retention of GA-rich mRNAs was a result of homeostatic regulation via R-MCD concentration, then the degree of nuclear retention should increase as the R-MCD expression increases over time. Using our PiggyBac cell line, we induced the expression of the reporter for 4, 8 and 12 hours, and sequenced the nuclear and cytoplasmic barcodes at each timepoint. We found that the nuclear retention of highly multivalent sequences was significantly stronger at 12 hours than at 4 or 8 hours (Figure 3F). We replicated this effect by transfecting the reporter plasmid pool for 8 or 24 hours and sequencing the nuclear and cytoplasmic barcodes (Figure S5B). In all experiments, the effect of sequence multivalency was greater than the effect of intron content, suggesting that the nuclear retention effect is not driven by changes in splicing (Figure S5A, S5C-D). We confirmed that the observed effects were not dependent on the stability of the reporter transcripts by expressing the pool for 16 hours, then measuring the relative abundance of transcripts over time after withdrawal of doxycycline, and found no relationship between the multivalency of transcripts and their stability (Figure S5G). Therefore, the expression level of the R-MCD dictates the selective retention of its own multivalent mRNA.

We next asked whether the expression of the reporter construct had a dose-dependent effect on the nuclear cytoplasmic localisation of endogenous mRNAs. As the time of dox induction resulted in a gradually increased concentration of mScarlet-PPIG_LCD_ (Figure 4A), we performed 3’ end sequencing on the nuclear and cytoplasmic fractions of our reporter cell line after 0, 4, 8 and 12 hours of doxycycline induction. The high quality of fractionation was stable across all samples (Figures S6A). We then categorised genes based on their total CDS GA-multivalency and looked at the nuclear-cytoplasmic distribution of these genes over time. The nuclear-cytoplasmic distribution of most mRNAs was unaffected by the expression of the reporter construct, but the group of mRNAs with high GA-rich multivalency became increasingly enriched in the nucleus over time, with a quantitative relationship between the degree of GA-rich multivalency and the degree of nuclear retention (Figure 4B, 4C). We performed a gene ontology analysis of the genes encoding mRNAs that were significantly more retained in the nucleus after PPIG_LCD_ expression, which identified enrichments in ontology terms related to chromatin, nuclear speckles, nucleolus and mRNA processing (Figure S6B). Significantly retained mRNAs encode proteins with a greater proportion of charged amino acids and high net-charge regions (Figure S6C, S6D). This demonstrates that increased abundance of a single R-MCD-protein leads to mutual homeostatic regulation of all charged proteins encoded by GA-rich mRNAs, a phenomenon we refer to as interstasis.

**Figure 4:**
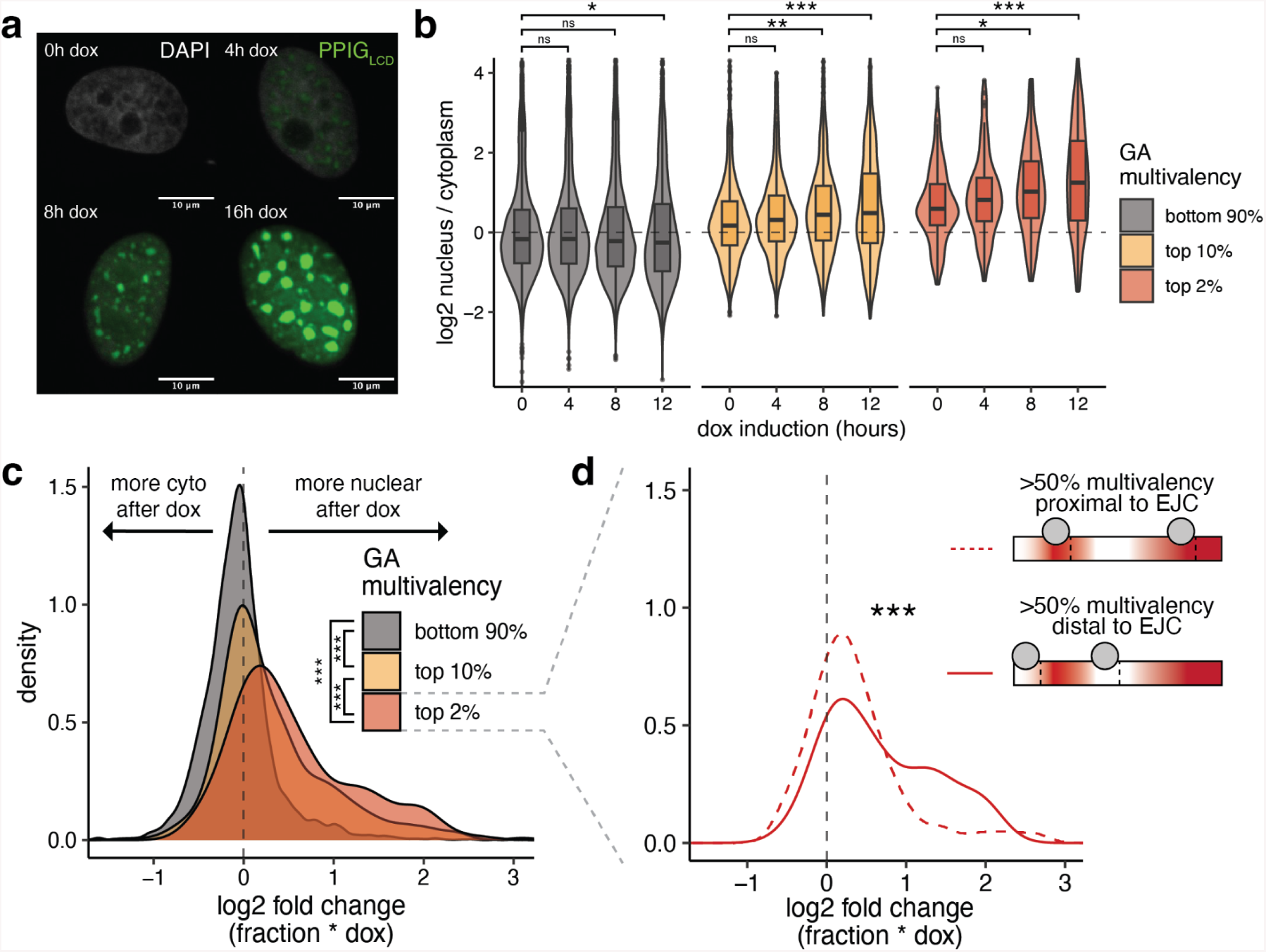
Interstasis of endogenous genes in response to R-MCD expression. a, Expression of PPIG_LCD_ over time after doxycycline induction. b, Nuclear-cytoplasmic distribution of mRNAs with different degrees of GA multivalency in response to different timepoints of PPIG_LCD_ expression. c, Per mRNA interaction effects describing the change in nuclear-cytoplasmic distribution after 12 hours of PPIG_LCD_ expression for mRNAs with different degrees of GA multivalency. d, Interaction effects from 4c for mRNAs in the top 2% of GA multivalency. mRNAs are classified based on whether 50% of the total purine multivalency occurs within 50 nt of a predicted exon junction complex site. All pairwise statistical comparisons performed using a Welch t-test with FDR correction for multiple testing, where * = p < 0.05, ** = P < 0.01, *** = P < 0.001.

To assess if the nuclear retention of GA-rich mRNAs decreases the synthesis of encoded proteins, we created codon-biased versions of a GA-rich region in *LUC7L3* mRNA that encodes an R-MCD, such that it either had a high or low GA-multivalency but equivalent GC content. We then cloned this region as a 3’UTR sequence downstream of a CDS encoding mGreenLantern. We transfected each of these two constructs into our mScarlet-PPIG_LCD_ reporter cell line, induced PPIG_LCD_ expression with doxycycline for 16 hours, and measured the intensity of mScarlet and mGreenLantern in each cell with flow cytometry. We found that as the expression of mScarlet-PPIG_LCD_ increased, expression of mGreenLantern strongly decreased when the 3’UTR of mGreenLantern contained a highly multivalent GA-rich sequence, whereas the effect was significantly weaker when the sequence was codon-biased to decrease the GA-content (Figure S6E). Interstasis thus enables a concerted response to increased concentration of an R-MCD protein by selective nuclear retention of GA-rich mRNAs, resulting in decreased synthesis of proteins encoded by these retained mRNAs and thereby maintaining the mutual homeostasis of R-MCD proteins.

### RNA features impacting interstasis

While the majority of mRNAs with high purine multivalency became more nuclear over time, the extent of this effect after 12 hours of reporter expression could be quite variable. Therefore, we examined additional features of the reporter, and of endogenous mRNAs, that affect the selectivity of their nuclear retention upon R-MCD expression (Figure 4C). Wherever an intron is spliced out of a nascent RNA, the exon junction complex (EJC) is deposited on the spliced mRNA, and the EJC then interacts with export factors such as ALYREF ^23, 24^, leading to ALYREF–EJC–RNA multimerization and packaging that facilitates loading of further mRNA packaging and export factors ^25^.

We were able to address the role of such splicing-mediated EJC loading with our reporter system, because the reporter pool was assembled such that each of the 8 fragments had a chance to contain an intron, and thus the total number of introns varied between reporters. If all 8 fragments included an intron, there were 8 exons between 162 and 211nt in length, and it was most common for reporters to have 4-5 introns. When the reporter pool was introduced into cells via plasmid transfection, the intron content had little influence on the nuclear-cytoplasmic distribution of spliced mRNA variants (Figure S5C, S5F). However, when the reporter pool was introduced by stable integration into the chromatin via a PiggyBac transposon system, the influence of intron content became more apparent, such that a greater number of introns resulted in more efficient nuclear export (Figure S5D). We categorised reporter genes based on their sequence multivalency and intron content and looked at their nuclear-cytoplasmic distributions. Highly multivalent transcripts became more abundant in the nucleus with increasing expression time when they had fewer introns, while multivalent transcripts with a large number of introns did not accumulate more in the nucleus over time (Figure S5E). This indicates that EJC-mediated packaging counteracts the nuclear retention driven by multivalent CDS sequences.

Given that the reporter system, when expressed from chromatin, identified an interplay between intron number and purine multivalency, we proceed to the analysis of endogenous mRNAs. We categorised mRNAs based on whether 50% of their purine multivalency was within 50 nt of an EJC. We found that mRNAs in which multivalent sequences were proximal to EJCs did not have a strong nuclear retention effect, but those in which the multivalent sequences were not proximal to EJCs were highly sensitive to PPIG_LCD_ expression (Figure 4D). Additionally, transcript length has also described to promote nuclear retention at steady state, and so we analysed the relative contributions of GA-rich multivalency and CDS length. We found in general, transcripts with longer CDSs are more retained in the nucleus after PPIG_LCD_ expression, but after correcting for length, GA-rich multivalency still promotes nuclear retention, and these two variables have a greater than additive interaction effect that drives stronger nuclear retention (Figure S6F). However, the change in nuclear-cytoplasmic localisation was only weakly predicted with exon density alone, and exon density and purine multivalency did not have a greater-than-additive effect on RNA localisation (Figure S6G). This suggests that EJC density alone may not influence the responsiveness of an RNA to R-MCD overexpression, but rather the placement of EJCs relative to multivalent RBP assemblies. Together, these findings indicate that GA-rich multivalency, transcript length and intron placement all impact the interstatic nuclear retention of GA-rich mRNAs, which could be explained by a competition between the EJC-driven mRNP packaging that promotes nuclear export, and speckle-localised RBPs that promote nuclear retention by binding to the GA-rich GeRM regions or to other parts of long mRNAs.

### Speckle proteins bind GA-rich RNAs

Many RNA binding proteins (RBPs) achieve high-affinity RNA binding through multivalent protein-RNA interactions, where similar motifs are concentrated along a stretch of RNA ^26^. This enables high-avidity binding of the whole motif cluster through multiple domains of the same RBP, or through multiple copies of RBPs that are usually stabilised through LCD-mediated interactions with each other^19, 27^. In this way, multivalent RNA sequences can also promote formation of higher-order protein-RNA condensates, which can be regulated by the concentration of RNAs and their bound RBPs. We therefore asked which RBPs preferentially bind to various types of GeRM regions in mRNAs, and which of these RBPs are most enriched on the GA-rich GeRM and thus contribute to the selectivity of nuclear export upon accumulation of R-MCD proteins.

We gathered public iCLIP and eCLIP datasets^28–32^ and ranked the RBPs based on the proportion of their crosslink events in the CDS, and their preference of crosslinking to their most-bound 5-mers in multivalent contexts (Figure 5A, B). For each RBP, we then compared the density of its crosslinking within each type of CDS GeRM region compared to the rest of the CDS of corresponding transcripts (Figure 5C). We observed that each type of GeRM region shows enriched binding of its own set of RBPs. The GA-rich multivalent regions were bound by a large number of RBPs that have been observed to be enriched in nuclear speckles (Figure 5C, bold)^28, 33, 34^, including six SR proteins that preferentially assemble on GA-rich GeRM regions, with the TRA2 SR proteins showing the strongest enrichment on these regions. For example, these SR proteins bind strongly and specifically to the GA-rich GeRM region that encodes the charged disordered domain of APP (Figure 5D). This raises the possibility that the speckle-localised SR proteins play a role in the interstasis of charged proteins.

**Figure 5:**
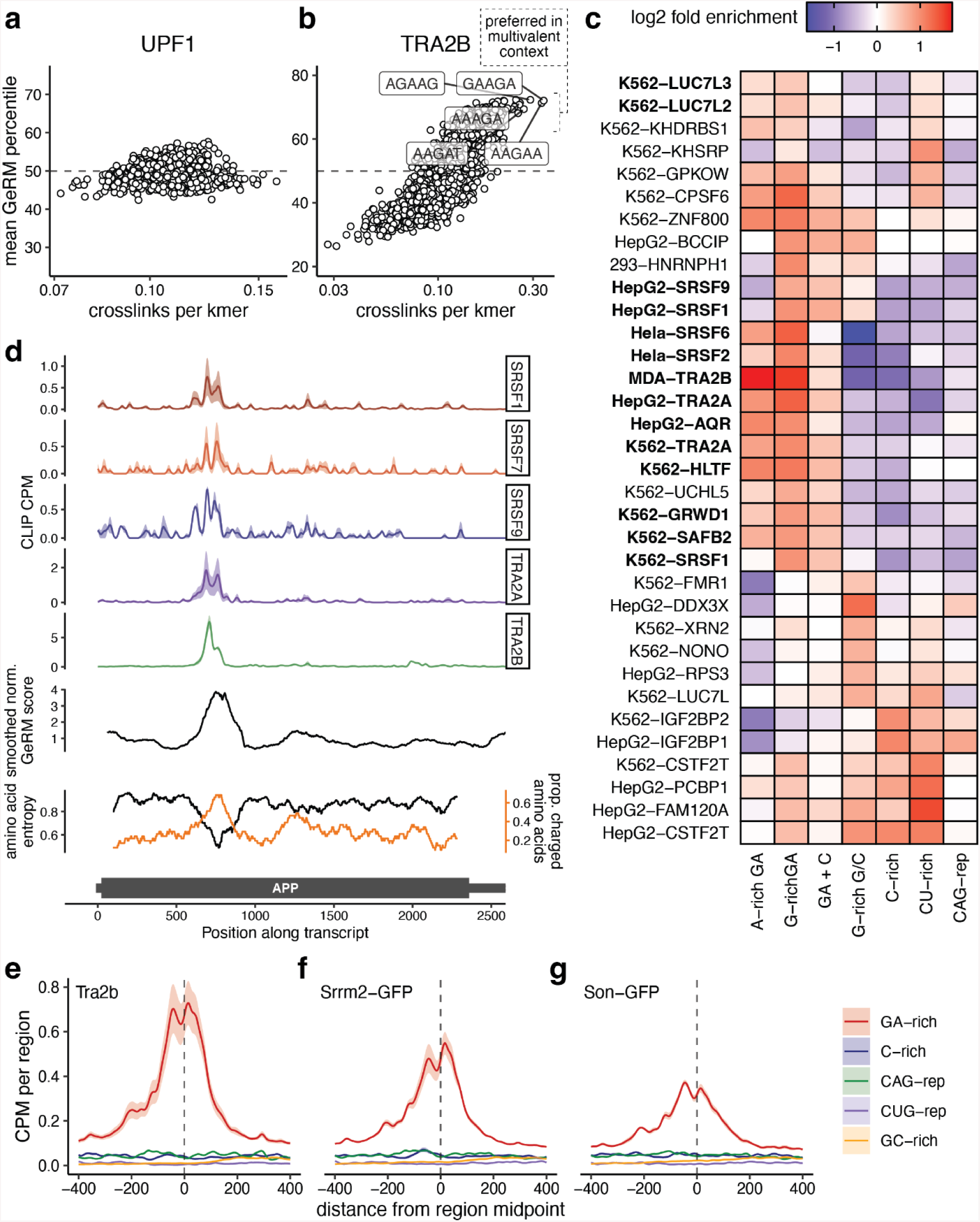
Nuclear speckle resident RBPs assemble multivalently on GA-rich multivalent regions. a-b, Plots depicting the CLIP crosslinking to each 5-mer and the mean percentile of the GeRM score of crosslinked 5-mers for UPF1 and TRA2B. c, A heatmap showing the fold enrichment for CLIP crosslinks falling within GeRM CDS regions compared to the rest of the transcript. d, Example CLIP crosslinking profiles for SR proteins across an APP transcript, with the smoothed GeRM score, amino acid entropy and proportion of charged amino acids. Solid lines represent the mean across two samples, while shaded regions represent the standard error. e-g, Metaprofiles of CLIP crosslinks for Tra2b, Srrm2-GFP (low molecular weight fraction) and Son-GFP (low molecular weight fraction) around the central points of subtypes of GeRM CDS clusters in mouse. Solid lines represent the mean across two samples, while shaded regions represent the standard error.

As SR proteins are well characterised to modify splicing, we investigated whether the splicing of exons with multivalent GA-rich sequences was perturbed by knockdowns of the proteins that bind them. We analysed public data in which both TRA2A and B, the strongest identified SR protein binders of GA-rich multivalent sequences, were knocked down with siRNA, which generally increases skipping of both alternative and constitutive exons ^29^. We found that exons with GA-rich GeRM regions were highly constitutive in both control and TRA2A/B double knockdown cells, with no systematic trend for their increased skipping upon the double knockdown (Figure S7A-B). Moreover, using data from VastDB ^35^, we found that GA-multivalent exons are highly constitutive across all available human tissues (Figure S7C). We conclude that strong binding of SR proteins to highly multivalent GA-rich exons is rarely associated with the modulation of alternative splicing.

To consolidate the findings from public data, we created two mouse embryonic stem cell (mESC) lines in which GFP tags were inserted at the N-termini of the core nuclear speckle proteins Son and Srrm2, and performed iCLIP for these GFP-tagged proteins using an anti-GFP antibody, as well as for Tra2b using an antibody against the endogenous protein. The SDS-PAGE analysis of Son and Srrm2 iCLIP showed strong signal of protein-RNA complexes below the molecular weights of Son or Srrm2, indicating that most RNA was derived from proteins smaller than Son and Srrm2 that were maintained in the immunoprecipitation despite high-salt and detergent washes (Figure S7D). The signal was not visible from GFP IPs in a wildtype cell line, therefore the signal was derived from proteins that interact with Son or Srrm2, likely because nuclear speckle proteins form strong RNP complexes that are resistant to high salt and detergents. Analysis of iCLIP crosslink profiles showed all three RBPs generally enriched on CDSs (Figure S7E), with the strongest enrichment around the GA-rich GeRM regions of CDSs (Figure 5E-G, S7G-H). Additionally, we found that iCLIP reads from all datasets were much more likely to contain a splice junction when the protein was bound to a GA-rich exon, confirming that these proteins are bound to multivalent sites post-splicing (Figure S7F). This confirms that Tra2b and core nuclear speckle-localised proteins are strongly bound to GA-rich GeRM regions in spliced mRNAs.

**Figure 7:**
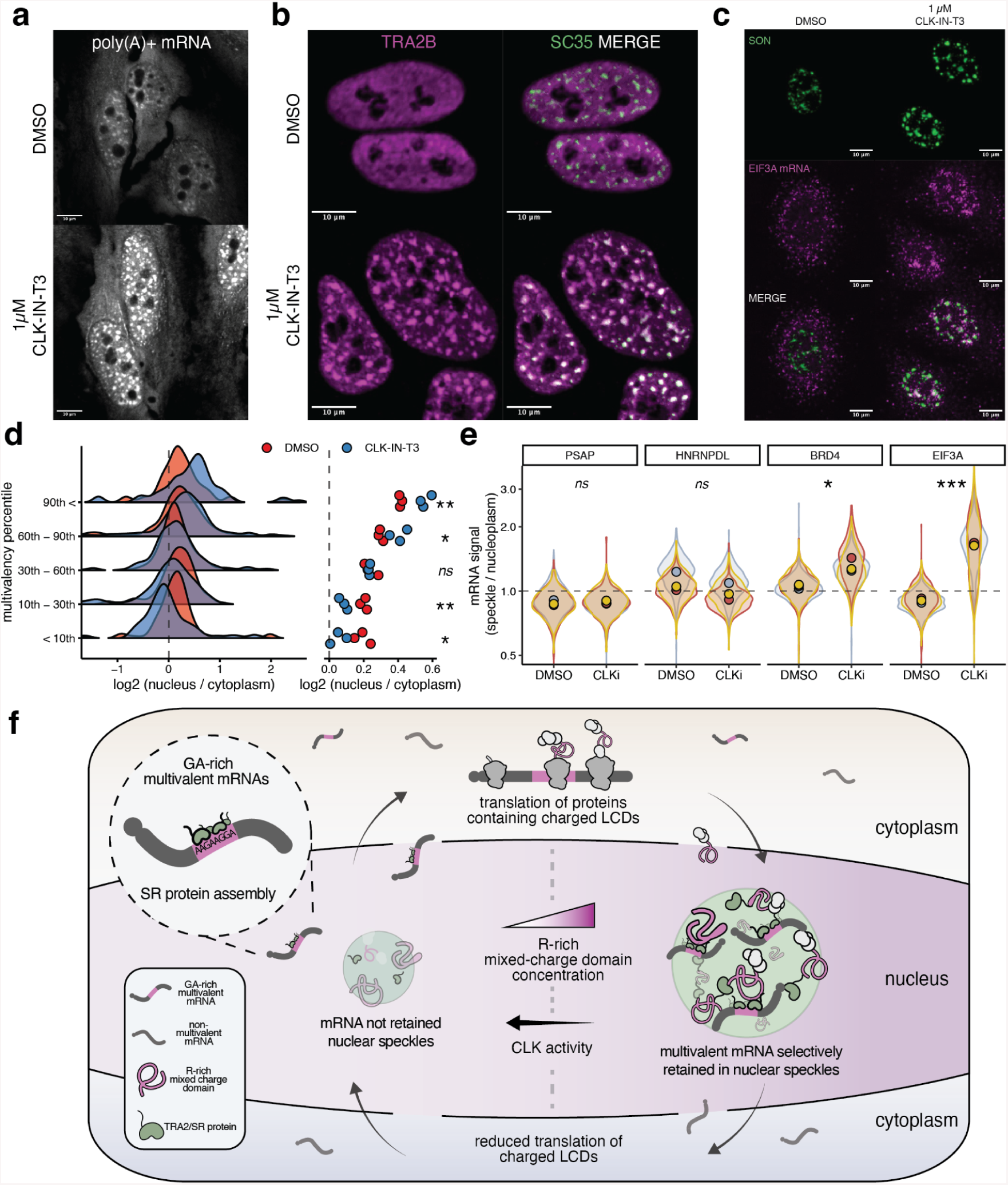
modulation of interstasis by CLK kinases. a, Oligo-dT FISH in cells treated either with DMSO or 1 µM CLK-IN-T3 for 8 hours. b, TRA2B and SC35 immunofluorescence in cells treated either with DMSO or 1 µM CLK-IN-T3 for 8 hours. c, Example image of SON immunofluorescence and EIF3A HCR-FISH in cells treated either with DMSO or 1 µM CLK-IN-T3 for 16 hours. d, The nuclear-cytoplasmic distribution of reporter transcripts binned by their total GeRM scores. Data from cells treated either with DMSO or 1 µM CLK-IN-T3 for 2 hours prior to induction of the reporter transcript pool for 6 hours. The right panel shows the group means per sample. e, Quantification of the enrichment of HCR-FSH signal within nuclear speckles versus nucleoplasm per nucleus per replicate for control (PSAP, HNRNPDL) or GA-multivalent (BRD4, EIF3A) mRNAs. f, A schematic detailing the interstasis of proteins with charged low complexity domains. All statistical comparisons performed using a Welch t-test with FDR correction for multiple testing, where * = p < 0.05, ** = P < 0.01, *** = P < 0.001.

### Nuclear speckle relocalisation marks interstasis

We asked if the speckle localisation of a GA-binding SR protein might be sensitive to the dose of R-MCD proteins, and thus could contribute to the GA-selectivity of the dose-dependent mRNA retention. We induced PPIG_LCD_ expression and performed immunofluorescence against TRA2B, the strongest identified binder of GA-rich GeRM regions. Indeed, TRA2B was diffuse throughout the nucleoplasm in the absence of PPIG_LCD_ expression, with only mild co-localisation with the nuclear speckle marker SC35 (Figure 6A). However, TRA2B became enriched in nuclear speckles 16 hours after the induction of PPIG_LCD_ (Figure 6A, 6B, S8A), which was dependent on the dose of PPIG_LCD_ (Figure 6B). We also observed a dose-dependent enrichment of poly-A(+) mRNA in nuclear speckles, accompanied by a mild increase in the proportion of nuclear mRNA, under these conditions (Figure 6C, S8B, S8C). Therefore, the retention of TRA2B and nuclear mRNA in speckles depended on the dose of R-MCD protein.

**Figure 6:**
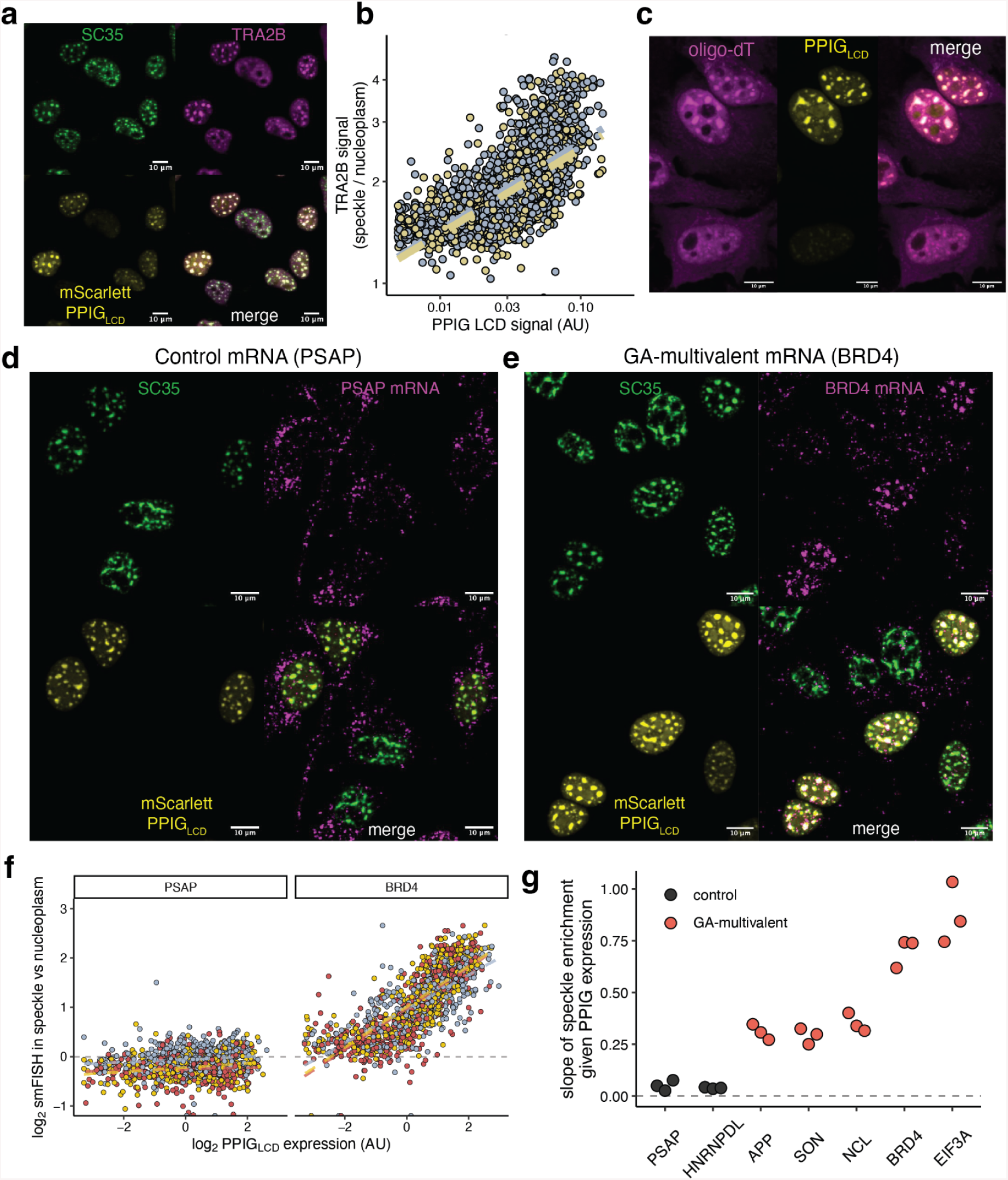
Dose-dependent sequestration of GA-multivalent mRNA and TRA2B in nuclear speckles. a, TRA2B and SC35 immunofluoresence in nuclei containing variable amounts of mScarlett-PPIG_LCD_. b, Quantification of the enrichment of TRA2B signal within nuclear speckles (based on SC35 signal) versus nucleoplasm per nucleus with respect to PPIG_LCD_ expression. Independent replicates are plotted in different colours and regression lines are plotted in dashed lines. c, Oligo-dT FISH of cells expressing mScarlet-PPIG_LCD_. d-e, Example images showing HCR-FISH signal for PSAP (control) or BRD4 (GA-multivalent) mRNAs, with SC35 immunofluorescence and mScarlett-PPIG_LCD_. f, Quantification of the enrichment of HCR-FISH signal within nuclear speckles versus nucleoplasm per nucleus with respect to PPIG_LCD_ expression. Independent replicates are plotted in different colours and regression lines are plotted in dashed lines. g, The slopes of linear regression models of the relationship between mScarlett-PPIG_LCD_ expression and mRNA enrichment within the speckle for control and GA-multivalent mRNAs. All statistical comparisons performed using a Welch t-test with FDR correction for multiple testing, where * = p < 0.05, ** = P < 0.01, *** = P < 0.001.

Induction of PPIG_LCD_ expression not only increases the expression of PPIG_LCD_ protein, but also the GA-rich mRNA that encodes it. To determine whether R-MCD proteins or the GA-rich mRNA itself drives the relocalisation of TRA2B to the nuclear speckle, we created a construct encoding mGreenLantern fused to the R-MCD of LUC7L3, and a construct in which a stop codon had been inserted upstream of the LUC7L3 R-MCD sequence, making it part of the 3’UTR. Two versions of each construct were made with opposing codon biases in LUC7L3, to create either GA-rich or GA-poor sequences (Figure S9A). We transfected HeLa cells with each of the four construct variants and performed TRA2B and SC35 immunofluorescence. When the LUC7L3 R-MCD was expressed, it localised to nuclear speckles (Figure S9C). We found that expression of the LUC7L3 R-MCD drove the speckle localisation of TRA2B independently of its GA-rich or GA-poor codon bias, whereas the effect was lost when a stop codon was inserted upstream of the LUC7L3 R-MCD sequence (Figure S9D, S9E). This demonstrates that TRA2B relocalisation and sequestration of GA-rich mRNA in the speckle is driven by the accumulation of R-MCD proteins, rather than the GA-rich mRNAs that encode them.

To assess the specificity of mRNA sequestration into nuclear speckles, we designed HCR-FISH probes against 5 GA-rich multivalent mRNAs and 2 non-multivalent control mRNAs. We then induced PPIG_LCD_ expression for 16 hours and performed HCR-FISH along with SC35 immunofluorescence (Figure 6D-E, S8D-E). We found a dose-dependent increase in the proportion of smFISH signal in the speckles relative to the nucleoplasm for all 5 multivalent RNAs, but not for either of the 2 control mRNAs, as evidenced by the slopes and fits of the regression models (Figure 6F-G, S8F-G). Thus, the selective retention of GA-rich multivalent mRNAs in speckles depended on the dose of R-MCD protein.

### CLK kinases modulate interstasis

Localisation of SR proteins within nuclear speckles is regulated by their phosphorylation via various kinases, such as the CLK family of kinases^36^. CLK kinase activity is regulated under a diverse array of physiological contexts, including during the cell cycle and in response to small variations in temperature^37, 38^. Given the potential role of SR proteins as mediators of interstasis, we asked whether the set point of interstasis could be modulated by the activity of CLK kinases.

We treated HeLa cells for 8 hours with 1 µM CLK-IN-T3, a selective inhibitor of CLK1, CLK2 and CLK3^39^, and observed a mild enrichment of poly-(A)+ mRNA and a strong relocalisation of TRA2B to nuclear speckles (Figure 7A-B, S10A-B). Upon 16 hours of CLK-IN-T3 treatment, the speckles became larger and fewer in number (Figure S10C-F). Thus, CLK kinase inhibition produces a similar phenotype to the overexpression of an R-MCD with regards to mRNA and TRA2B localisation and nuclear speckle morphology.

To test the selectivity of speckle mRNA localisation, we treated HeLa cells with 1 µM CLK-IN-T3 for 16 hours and repeated HCR-FISH against the GA-rich EIF3A and BRD4 mRNAs as well as the control mRNAs PSAP and HNRNPDL. CLK inhibition significantly increased the localisation of EIF3A and BRD4, but not control mRNAs, to nuclear speckles relative to the nucleoplasm (Figure 7C, 7E, S10D-E). We confirmed that CLK inhibition drives nuclear retention of mRNAs using our reporter system. We treated cells with 1 µM CLK-IN-T3 or DMSO for 2 hours, then induced the expression of the reporter transcript pool with doxycycline for 6 hours, at which point the expression of the PPIG_LCD_ protein is still negligible, maintaining the drug treatment during this time. We found that CLK inhibition significantly increased the nuclear retention of the most multivalent reporter transcripts, and relatively decreased the nuclear retention of the least multivalent reporter transcripts (Figure 7D).

To more globally assess the changes in nuclear-cytoplasmic distribution of mRNAs, we then treated HeLa cells with 1 µM CLK-IN-T3 for 8 hours, collected nuclear and cytoplasmic fractions and performed 3’ end sequencing from both fractions. We found that mRNAs with GA-rich GeRM regions in CDSs were significantly more retained in the nucleus following treatment with CLK-IN-T3 (Figure S10G). The extent of nuclear retention of each transcript was well correlated when comparing the effects of 8 hour CLK inhibition and 12 hour PPIG_LCD_ expression (Figure S10H). These results further suggest that nuclear retention of GA-rich mRNAs results from the alteration of the properties of the nuclear speckle, including increased speckle localisation of TRA2B, which can be induced either by increased abundance of R-MCD proteins, or by decreased phosphorylation of SR proteins.

## Discussion

The dosage of many LCD-containing proteins is critical for their function and to maintain cellular health^1, 2, 40^. As proteins with similar LCDs often localise to the same condensate, where they often function together (with nucleolus and nuclear speckles as examples)^4^, and thus it would be advantageous for their collective homeostasis to be maintained mutually. Here we identify interstasis, a mechanism of such mutual homeostasis by which the dosage of related LCDs is controlled as a group.

Through a new computational approach, we quantified the multivalency potential of RNA sequences. We find that distinct classes of multivalent CDS regions encode specific types of LCDs in functionally related sets of proteins. Notably, the CDS regions exhibit conserved codon biases that promote their multivalent identity. Focusing on the GA-rich multivalent CDS regions that encode charged LCDs, we developed a new combinatorially-assembled reporter strategy to demonstrate the role of these codon biases in interstasis. We show a dose-response relationship between R-MCD expression and sequestration of GA-rich multivalent mRNA in the nuclear speckle, resulting in decreased mRNA export and reduced translation of proteins containing charged LCDs (Figure 7F). This retention is mediated by RBPs, especially SR proteins such as TRA2B, which assemble along GA-rich multivalent sites. Additionally, we show that inhibition of the CLK family of kinases phenocopies R-MCD expression, likely by modifying the interactions between SR domains and other speckle components.

Therefore, multivalent protein-RNA interactions exploit the structure of the genetic code to specifically assemble on regions in mRNAs that encode functionally related LCDs. This allows the cell to modulate the fate of that mRNA in response to existing LCD abundance or cellular signalling. Previous work has shown that the abundance of R-MCDs alters the condensation properties of the nuclear speckle^8^. Biomolecular condensates are commonly conceptualised as percolated interaction networks of multivalent molecules^41, 42^. Our work suggests that participation of multivalent mRNAs and their binding partners in this interaction network is dose-sensitive and tunable by signalling, which allows direct feedback from the biophysical properties of the condensate onto the mRNAs that encode those properties.

Several sequence motifs and RBPs are known to alter the export of mRNA at steady state^22, 43, 44^. We now show that mRNA export is also leveraged to regulate a specific group of mRNAs as a mechanism for maintaining the homeostasis of functionally related proteins. We show that hypophosphorylation of SR proteins such as TRA2B promotes GA-multivalent mRNA localisation within the nuclear speckle, while EJC-mediated mRNP packaging competes with SR proteins to reduce speckle localisation. Many RBPs are constitutively enriched in nuclear speckles, so the valency of an mRNA scales with its length in addition to the local multivalent regions. Other mechanisms likely exist to recruit specific sets of RNAs to the speckle, or to promote the global retention of mRNAs within the speckle^45^.

We find that the activity of CLK family kinases can modulate interstasis of charged proteins via GA-rich mRNAs. CLK signalling is responsive to a great variety of physiological states, including cell cycle progression and small variations in temperature^37, 38^, indicating a broad capacity to adjust the threshold of interstasis depending on the cellular context. The role of CLKs in interstasis might be relevant to their implication in various diseases and consideration as potential therapeutic targets^36^. Interestingly, the multivalency-enhancing bias in arginine codons is strongest in R-LCD-encoding sequences of vertebrate clades that evolved on land (amniotes), and is also more pronounced in the non-vertebrates that live on land, which generally face the challenge to balance cellular O_2_ delivery against oxidative damage^46^. Notably, highly charged proteins are at particular risk of oxidative damage, which can lead to their unfolding and aggregation in ageing cells^47^. Indeed, proteins containing mixed charge domains have been observed to be highly aggregated in the brains of Alzheimer’s disease patients^48^. Given that protein dosage control is key to avoiding aggregation, interstasis of charged proteins may be important for minimising the accumulation of oxidative damage during ageing.

Beyond mechanisms for interstasis of charged proteins, our discovery of conserved multivalent codon biases within LCD-encoding regions could prove valuable for modelling and interpreting the impact of synonymous mutations on human diseases^49^, and for the design of synonymous gene-recoded therapeutics^50^. Moreover, analysis of public CLIP data showed that each class of multivalent RNA regions is preferentially bound by a distinct set of RBPs. We find that other types of multivalent CDS regions contain strong codon biases that reinforce multivalency, and tend to encode functionally coherent sets of proteins. It remains to be tested whether the total abundance of other types of LCD-containing proteins might also be controlled through interstasis, which could involve various condensates that attract each type of LCD in a dosage-sensitive manner, enabling feedback regulation of the multivalent mRNAs that encode a type of LCD-containing proteins. Since many age-related diseases are caused by low-complexity repeat expansions^51^, or aberrant phase separation and aggregation of LCD-containing proteins^5^, and many other diseases involve disruptions in nuclear speckles^52^, it will be important to assess whether interstasis and its perturbation plays a role in these diseases.

## Supporting information

Supplemental Table 1 (Species)

Supplemental Table 2 (Reporter)

Supplemental Table 3 (Targeted Seq)

Supplemental Table 4 (3' End Seq)

Supplemental Table 5 (LUC7L3 Sequences)

## Acknowledgments

The authors would like to thank the following Francis Crick Institute Science Technology Platforms for their technical support: Advanced Sequencing, Advanced Light Microscopy, Flow Cytometry and Cell Services. The authors also thank the BioOptics Facility at the Institute of Molecular Pathology. This work was supported by the European Research Council under the European Union’s Horizon 2020 research and innovation programme (835300-RNPdynamics) and Wellcome Trust Investigator Award (215593/Z/19/Z) to J.U., by the Francis Crick Institute which receives its core funding from Cancer Research UK (CC0102), the UK Medical Research Council (CC0102), and the Wellcome Trust (CC0102) and by the UK Dementia Research Institute [award number UK DRI-RE13553 and RE21605] which receives its funding from UK DRI Ltd, funded by the UK Medical Research Council, Alzheimer’s Society and Alzheimer’s Research UK. For the purpose of Open Access, the author has applied a CC BY public copyright licence to any Author Accepted Manuscript version arising from this submission.

## Author Contributions

R.F. and J.U. conceived and jointly supervised the study. R.F. conducted data analysis, designed and performed experiments. N.C.H., H.D. and L.K. also performed experiments. A.M.C. and I.A.J. assisted with the coding of the GeRM package. O.G.W. assisted with the design of the reporter construct pool. C.P. and S.L.A. provided additional experimental designs. R.F. and J.U. wrote the manuscript, with contributions from all co-authors. All authors agreed with the content and consent to submit the manuscript with their contributions.

## Methods

### GeRM algorithm

GeRM is calculated from a string of consecutive overlapping nucleotide sequences of length 𝑘 (k-mers). From the set of 𝑛 consecutive k-mers 𝐴, the GeRM score 𝑔 of the central k-mer 𝐴_0_ is calculated as follows:

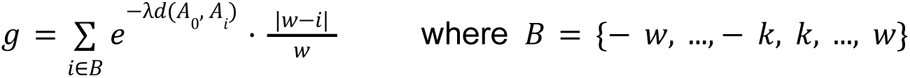

where 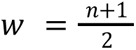, the function 𝑑 represents Hamming distance, and λ represents a scaling constant. In non-mathematical terms, the GeRM score is calculated by comparing a k-mer to all the other k-mers that surround it in a fixed window. For each of the surrounding k-mers, the sequence similarity to the central k-mer is calculated from the negative exponent of the Hamming distance, such that k-mers with identical sequences have a high score and those with unrelated sequences have a low score. The constant λ determines how quickly this similarity score decays as sequences become more dissimilar to the central k-mer. This sequence similarity score is multiplied by a distance score, which decays linearly from 1 to 0 with distance from the central k-mer. k-mers that overlap with the central k-mer are ignored. For k-mers at the edges of transcripts, where the window exceeds the end of the transcript, all positions that fall outside of the transcript are given a score of 0. The sum of all the distance-weighted sequence similarities is summed to give the GeRM score.

In this manuscript, we used a k-mer length (𝑘) of 5, a window size (𝑛) of 123, and a scaling factor (λ) of 1. We normalised the GeRM scores such that the minimum value in the transcriptome was 0 and the median was 1. Smoothing of GeRM scores was performed by taking the mean of values in a sliding window of size 123.

For the calculation of codon-shuffled multivalency, all codons for a given amino acid in each transcript were swapped such that the number of instances of each codon was preserved for each transcript, but their order was randomised. All GeRM scores for each transcript were calculated, and the mean GeRM score per position across ten shuffles was calculated.

The GeRM algorithm is available as an R package from https://github.com/ulelab/germ.

### Identification and classification of GeRM regions

For each protein-coding gene in GENCODE 29 or GENCODE M22, the transcript with the longest CDS was selected and ties were broken by the longest total transcript length. Only spliced transcripts were used for analysis. CDS GeRM regions were defined by taking all positions where the smoothed GeRM scores exceeded the 98^th^ percentile of smoothed GeRM scores within the CDS, and GeRM regions were adjusted to contain all k-mers that fell within the smoothing window. GeRM regions with at least 41 nt of overlap were merged. GeRM regions for which at least ⅓ of the region fell within an untranslated region were excluded.

For each GeRM region, the relative proportion of the region’s total multivalency that was accounted for by each possible k-mer was calculated. For clustering, UMAP was used to reduce the dimensionality of the data to 4 dimensions, using 50 neighbors and a minimum distance of 0.001. The reduced data was clustered first using OPTICS with the minimum points defined by 1% of the total number of GeRM regions. Clusters were then obtained using DBSCAN. The reachability threshold for DBSCAN was defined by determining how the proportion of points below a given threshold changed as the reachability threshold decreased (from the 99th percentile to 0), then taking the point at which the first derivative was 40% of the maximum (the knee of the plot). GeRM regions falling outside of a cluster were excluded from further analysis. For presentation purposes, UMAP was used to reduce the original data to have two dimensions, with other parameters kept the same.

### Low complexity and disorder

To define the complexity of amino acid sequences, information entropy was calculated using the R package HDMD in a sliding window of 41 amino acids along the translated CDS sequences from the set of transcripts used for GeRM scoring. Low complexity domains (LCDs) were defined as regions with entropy in the bottom 2% of all entropy scores, with all amino acids in the sliding window being considered part of the LCD. LCDs that overlap were merged. AlphaFold^53^ pLDDT scores were obtained for the same set of proteins where available from UniProt.

### GeRM codon ratios

To determine whether a given position in a sequence supported the local multivalency of that sequence, we defined a GeRM codon ratio. First, ‘mutable positions’ were defined as any position that could be synonymously mutated. Primarily, these are the final position of the codon, but the first positions of some leucine and arginine codons can also be synonymously mutated. For each mutable position, sequences were produced in which every possible synonymous mutation was made. The GeRM scores of all 5-mers that overlapped with the mutable position were calculated, and the maximum of these GeRM scores was taken for each possible mutation or the native sequence. Then, the maximum GeRM score associated with the 5-mers overlapping the native sequence was divided by the mean of the maximum GeRM scores associated with all possible mutations to create a ratio. When this ratio is positive, the native nucleotide at the mutable position is associated with higher GeRM than the other possible mutations that could exist at that position on average.

### Conservation of GeRM

PhyloP conservation across 100 vertebrates and 470 mammals in hg38 coordinates were downloaded from the UCSC Golden Path. These values were converted into transcriptomic coordinates based on the single transcripts per genes that were used for GeRM calculation. Mutable positions were binned based on their GeRM codon ratio with the binning calculations being applied to each codon separately, such that each bin contained an equal proportion of each codon.

For raw conservation analysis, the average PhyloP conservation values of mutable positions in each GeRm codon ratio bin were used. For analysis of normalised conservation values, the PhyloP of mutable positions were normalised for each codon by subtracting the median conservation of that mutable position in that codon by the median conservation across all mutable positions of the same type. For example, the conservation value associated with one instance of the A in the codon CGA would be normalised by subtracting the median conservation of all As in CGA codons. Each normalised position was then further normalised by subtracting the PhyloP value of the middle position of the codon (in the previous example, the G in that specific instance of the CGA codon). Subtraction was used for normalisation because PhyloP scores operate on a logarithmic scale.

### Arginine codon usage

For analysis of arginine codon usage in R-LCDs, all human LCDs were filtered for those that obtain at least 20% arginine residues. The proportion of each arginine codon in the respective coding regions was calculated. Pairwise correlations between arginine codon usages using Spearman’s rank correlation. To define CG- and GA-rich codon groups, principal component analysis was performed based on the arginine codon usage for all R-LCDs, and the top and bottom third of LCDs based on the first principle component were defined as CG- and GA-rich R-LCDs respectively.

For the definition of R-LCDs in different species, all CDS for each species were downloaded from Ensembl and a single transcript per gene was identified according to the previously described approach. LCDs encoded by these sequences were defined as previously described, using the complexity threshold defined for the human proteome for each species. A full list of the species and annotation versions can be found in Supplemental Table 1.

### Gene ontology

Gene ontology analyses were conducted using the topGO R package and using the weight01 algorithm to account for the topology of the gene ontology graph.

### Reporter library design

The mScarlet-PPIG_LCD_ library was designed by concatenating the coding sequences for mScarlet and PPIG_402-684_. The concatenated sequence was then iteratively synonymously mutated such that the GC content of the sequence was maintained within 2% of the original sequence, choosing codons with fewer As and Gs until the number of AG-only 5-mers reached a minimum. Untranslated regions were added to this sequence. Seven equally spaced sequences of 28 nt were chosen along the sequence to be used as overlap sequences for future Gibson assembly. These overlap sequences were protected from future *in silico* mutagenesis.

26 constitutively spliced introns ranging from 83 nt to 384 nt in length were selected from housekeeping genes. We generated 25 different combinations by randomly choosing 8 of these introns, then performed the following iterative optimisation algorithm to each of these 25 combinations to generate sequences with optimised predicted splicing.

First, introns were inserted in the reporter coding sequence in positions that were exactly between the previously defined overlap sequences or the start and end codons. The probability of donor and acceptor site usage was predicted using SpliceAI, and all probabilities were summed. Next, 100 coding sequences with 5 random synonymous mutations were generated and a sequence was selected that best maintained GC content and kept GA 5-mer content low. After this, an intronic sequence was selected with a 80% chance to be mutated at 5% of its positions and a 50% chance to be shifted by up to 20 nt. The probability of donor and acceptor site usage in the new sequence was predicted with SpliceAI, and the sum of the probabilities was compared to the previous sequence. If the total probability of splice site usage was increased in the mutated sequence, then the mutated sequence was put forward for another round of mutation, until 2000 iterations were completed.

To create a GA-rich version of the reporter coding sequence, the iterative process was repeated, but with synonymous mutations that maintained GC content but maximised GA-rich 5-mer content. The sequence and position of the introns was kept constant for this process.

To determine which of the 25 possible combinations of 8 introns resulted in the best predicted splicing in both GA-poor and GA-rich reporter sequences, we selected the intron set with the highest summed probability of predicted donor and acceptor site usage. This resulted in GA-poor and GA-rich codon-biased reporter sequences in which both of the splice sites for each of the 8 introns was predicted to have at least a 98.5% probability of being spliced.

For both the GA-rich and GA-poor sequence, 8 sequence segments were generated that spanned from each overlap sequence to the next (or to the untranslated region) and that either contained an intronic sequence or not. Therefore, each of the 8 segments of the final reporter gene varied in two possible ways: GA-richness and the presence or absence of an intron, and each of these variable segments shared a fixed overlap sequence with the neighboring segment.

### Reporter library generation

All 32 possible segment variations were ordered as dsDNA fragments from IDT. In addition to the overlap sequences that would be part of the final reporter transcript, additional overlap sequences were added for intermediate assembly steps. Specifically, an overlap sequence was added to the 3’ end of the first segment and to the 5’ of the last segment, and another overlap sequence was added to the 3’ ends of the first and fifth segments and the 5’ ends of the fourth and last segment. Then, two pools containing all variations of the first four segments or all variations of the last four segments were made, such that intron-containing variants had twice the molarity of intronless variants at each position, but all positions had the same overall molarity. A Gibson assembly reaction was carried out on each pool, and due to the additional overlaps on the first, fourth, fifth, and final segments, circular pieces of dsDNA were formed, each containing one of the four variations at each of the four positions used for that pool.

These circular pieces of DNA were then linearised and amplified by PCR, and the linearised fragments deriving from the first and second half of the reporter gene were again Gibson assembled together. Again, because of the additional overlap sequences attached to the first and last segments, the resulting assemblies were circular. Each assembly contained a 5’UTR, a coding sequence encoding mScarlet-PPIG_LCD_ but with a variable amount of GA-rich 5-mers and a variable number of introns, and a 3’ UTR. These assemblies were then linearised and amplified by PCR and Gibson assembled into a plasmid backbone (pTwist-CMV) containing a CMV promoter and a barcoded 3’UTR sequence with a poly-adenylation site. Therefore, the plasmids in the pool contained different versions of the reporter gene, each with a unique barcode in their 3’UTR.

After each reaction, the product was purified using Mag-Bind TotalPure beads. All PCRs were performed using Q5 High-Fidelity Master Mix (NEB). All Gibson assemblies were performed using NEBuilder HiFi DNA Assembly (NEB).

To generate a dox-inducible PiggyBac integration vector containing the reporter library (ePB-ExportReporter), reporter genes and their barcoded UTRs were amplified out of the original reporter plasmid library with a low number of PCR cycles, before being inserted into an all-in-one PiggyBac plasmid (ePB-Bsd-TT-NIL) containing a doxycycline-inducible promoter as well as a constitutive promoter driving the expression of the rtTA protein and blasticidin resistance gene. All designed fragments, as well as example reporter sequences and plasmids, are provided in Supplemental Table 2.

### RNA-extraction

RNA extractions were performed from cell pellets using the Maxwell RSC simplyRNA kit (Promega) using a Maxwell RSC Instrument (Promega).

### Long-read sequencing

For sequencing of the plasmid pool, the reporter genes were amplified from the plasmid DNA with a minimal number of PCR cycles. For sequencing of the reporter RNA, one well of a six well plate of Hela cells were transfected with 500 ng of the reporter plasmid pool and 2000 ng of a filler pUC19 plasmid for 16 hours. RNA was extracted and 500 ng of total RNA was used as input for a SuperScript IV reverse transcription (RT) (Invitrogen) reaction using an oligo-dT reverse transcription following manufacturer instructions. The reaction was purified with magnetic beads, then amplified according to the same approach as for the plasmid DNA. The specificity of the amplicon was confirmed with an agarose gel. In both cases, the PCR product was purified with magnetic beads (Mag-bind TotalPure NGS; Omega-Bio-tek) and Nanopore adapters were added using the Nanopore Ligation Sequencing Kit (Oxford Nanopore according to manufacturer instructions. The libraries were sequenced separately using MinION R9.4.1 Flow Cells (Oxford Nanopore).

Raw Nanopore data was basecalled with Guppy 6.0.1. The identity of the segments in each plasmid sequence or transcript sequence were determined by fuzzy string matching using the Python package rapidfuzz, and the plasmid sequencing data was used to relate the plasmid identities to their 3’UTR barcodes.

### Targeted sequencing

Targeted sequencing libraries were prepared from 500 ng of RNA input per sample. A targeted reverse transcription was performed with SuperScript IV (Invitrogen) using an RT primer which was specific to the reporter 3’UTR and contained a 4nt experimental barcode, a 10nt UMI and an Illumina p7 adapter sequence. The reaction was performed according to manufacturer instructions. Samples were multiplexed in groups of 6 at this point, and the RNA was removed by alkaline hydrolysis for 15 minutes, the pH was neutralised with hydrochloric acid and the cDNA was purified with magnetic beads. A nested PCR was then performed in which 10 cycles with one set of primers was performed. 5% of the reaction was spiked into the next 6 cycle reaction, in which an Illumina p5 adapter sequence was added. The PCR reaction was purified with magnetic beads (Mag-bind TotalPure NGS; Omega-Bio-tek) and Illumina i5 and i7 indices were added by a final PCR. Libraries were sequenced on a NovaSeq 6000 by the Advanced Sequencing Facility at The Francis Crick Institute. All primer sequences can be found in Supplemental Table 2.

### Analysis of targeted sequencing

Targeted sequencing libraries were analysed in R. Libraries were demultiplexed using the barcodes introduced by the RT primer, and the plasmid barcode was extracted by matching the flanking constant regions using fuzzy string matching using the R package stringdist. Reads containing the same barcode were deduplicated by their UMI sequences. Barcodes were counted and related to their reporter gene characteristics based on the long read sequencing data. Barcode counts were normalised to the total number of assigned barcodes for each sample. Barcodes with low numbers of counts were excluded from further analysis. Nuclear-cytoplasmic ratios for each barcode were calculated for each replicate. All processed data for each experiment can be found in Supplemental Table 3.

### 3’ end sequencing

For each sample, 250 ng/µL of RNA was fragmented in 10 mM Tris-HCl pH 7.5, 10 mM MgCl2 buffer for 5 min at 95°C. 2 µL of fragmented RNA, 2 µL water, 0.5 µL of 10 mM dNTPs and 0.5 µL of 5 µM of oligo-dT reverse transcription primers containing Illumina P7 sequences were mixed and primers were annealed by heating at 65°C for 3 min then cooling to 42°C at 1°C/s. At this point, reverse transcription was carried out using SuperScript IV (Invitrogen) according to manufacturer instructions, with the addition of 0.25 µL of 40 µM of a template-switching oligo containing Illumina P5 sequences and UMIs. The reaction was incubated for 1 hour at 42°C. Following alkaline hydrolysis of RNA, the product was purified using magnetic SPRI beads and amplified using 0.5 µM i5/i7 Illumina indexed primers in Q5 High-Fidelity Master Mix (NEB).

### Analysis of RNA-seq data

All sequencing data was mapped to the GRCh38.p12 genome using the GENCODE v29 basic genome annotation. All 3’ end sequencing data was trimmed using cutadapt v4.4^54^ to remove poly-A sequences and Illumina adapters. All data was processed using the nf-core RNA-seq Nextflow pipeline v3.12.0^55^, and in the case of the 3’ end sequencing data, the additional option --noLengthCorrection was provided to Salmon to prevent length correction for gene expression.

Differential expression analysis was conducted using DESeq2 in R^56^, with effect size shrinkage using the apeglm package^57^. Changes in nuclear-cytoplasmic ratio between two conditions were assessed using the interaction term of the models (∼ fraction * time). Splicing analysis was conducted using rMATS v4.1.2^58^ using the skipped exons calculated using the JCEC quantifications.

To classify genes based on their degree of purine multivalency, a purine multivalency score was calculated for each gene based on the coding sequence of its primary transcript as previously defined. First, for each of the 3 previously identified GA-rich GeRM clusters, the proportion of total multivalency that was accounted for by each 5-mer was calculated, and 5-mers were sorted in descending order based on this proportion. The cumulative sum was calculated and all 5-mers that accounted for the first 50% of the multivalency in each GeRM region were taken forward as representative 5-mers for those regions. Then, 5-mers were scaled to all other instances of the same 5-mer, such that their mean score was 0 and their standard deviation was 1. For each transcript, the sum of all of these scaled GeRM scores was summed across all representative 5-mers. Based on this metric, transcripts that have many instances of GA-rich 5-mers specifically in multivalent contexts are rewarded. All processed data from 3’ end sequencing experiments can be found in Supplemental Table 4.

### Immunofluorescence

Immunofluorescence was performed by fixing cells in 4% PFA and 0.1% glyoxal at room temperature for 10 minutes. Samples were permeabilised by incubating with PBS with 0.5% Triton-X for 5 minutes at room temperature. Samples were blocked using PBS with 3% BSA and 0.1% Tween (blocking solution). Primary antibody incubations were carried out in blocking solution for 1 hour at room temperature or overnight at 4°C. Samples were washed 3 times with PBS with 0.1% Tween for 5 minutes, and secondary antibody incubations were carried out in blocking solution for 1 hour at room temperature in the dark. Samples were washed twice, then incubated with 200 ng/mL DAPI for 15 minutes in the dark at room temperature. DAPI was washed out and samples were placed in glycerol mounting media (90% glycerol, 20 mM Tris pH 8, PBS).

The following antibodies were used at the following concentrations: rabbit anti-TRA2B (1:1000, abcam, ab31353), rabbit anti-SON (1:1000, Sigma, HPA023535), mouse anti-SC35 (1:500, Sigma, S4045). Secondary antibodies against mouse and rabbit, conjugated to AlexaFluor-488 or AlexaFluor-647 were used at 1:500 dilution (Thermo).

### HCR-FISH

For each mRNA, HCR 12 probe pairs were designed with the B4 amplifier sequences using the HCR 3.0 Probe Maker^59^. HCR 3.0 was performed on cells in µ-Slide 8 Well Glass Bottom slides (Ibidi, 80827) as described in the original protocol^60^, with the following modifications. First, fixation and permeabilisation was performed as described in the immunofluorescence section. After pre-hybridisation, primary probe hybridisation was performed with a probe concentration of 10 nM for 3 hours. Overnight HCR amplification was performed in a volume of 125 µl with half the concentration of HCR hairpins (Molecular Instruments). Following HCR amplification, samples were washed according to the original protocol, and then immunofluorescence was performed as described previously, but with all buffers containing 2XSSC in place of PBS.

### oligo-d(T) FISH

Samples for oligo-d(T) FISH were fixed as described for immunofluorescence. Samples were incubated with HCR 3.0 hybridisation buffer for 30 minutes at 37°C, then in hybridisation buffer containing 1 µg/mL oligo-dT(25)-Cy5 for 2 hours. Samples were washed twice in 5XSSC with 0.1% Tween, stained with 200 ng/mL DAPI in the same buffer and washed once more before glycerol mounting media was added.

### Microscopy

Microscope images were acquired using a Olympus IX3 Series (IX83) inverted microscope, equipped with a Yokogawa W1 spinning disk and a Hamamatsu Orca Fusion CMOS camera (pixel size: 6.5μm, 2304×2304, 5.3 megapixels).

### Analysis of microscopy data

All images were z-projected using a maximum projection. Segmentation of nuclei and cytoplasms was performed using Cellpose v2.0^61^ using the cyto2 model. Segmentation of nuclear speckles was performed using CellProfiler using the Otsu thresholding algorithm, and nucleoplasms were considered all non-speckle regions of the nucleus. Mean intensities in the nucleoplasm and nuclear speckle were calculated per cell, but statistical comparisons between groups counted each replicate as a single observation, taking the mean values across all cells in the replicate.

### Flow cytometry

For each sample, a single well of a 6-well plate of ePB-ER HeLa cells was transfected with one of the mGreenLantern LUC7L3-LCD reporter plasmids in which the LUC7L3 sequence was used as a 3’UTR but was either GA-rich or GA-poor. Doxycycline was added to the media at the point of transfection. Four replicates were performed for each plasmid, as well as untransfected and uninduced controls. After 16 hours, the cells were dissociated, washed twice with ice cold PBS and kept on ice while the samples were measured with an LSR Fortessa (BD Biosciences). The intensities of mGreenLantern and mScarlet were measured using a 488 nm Blue Laser (Coherent Sapphire 100 mW) with a 530/30 nm filter, and a 561 nm Yellow Green Laser (Coherent Sapphire 100mW) with a 610/20 nm filter, respectively.

Gating of singlets was performed using FlowJo, and subsequent analysis was performed in R. For each plasmid, the mGreenLantern intensities were normalised to the mean intensity values of the uninduced control samples. Cells were binned according to their mScarlet intensity values, and the mean mGreenLantern intensity values for all cells in each bin was calculated per replicate.

### Plasmid construction

Double stranded DNA fragments were ordered as gBlocks or eBlocks from IDT. Primers were ordered from Sigma-Aldrich. All PCRs utilised Q5 High-Fidelity Master Mix (NEB) or Phusion High-Fidelity Master Mix (NEB). Backbone plasmids were linearised via PCR followed by treatment with DpnI (NEB) before assembly with fragments using NEBuilder HiFi DNA Assembly (NEB) according to the manufacturer’s protocol. Phosphorylations and ligations were performed using T4 PNK kinase (NEB) and T4 DNA ligase (NEB). Assembly reactions were purified using magnetic SPRI beads (Mag-bind TotalPure NGS; Omega-Bio-tek) before DNA transformation into 5-alpha Competent *E. coli* (NEB). All sequences were confirmed via Sanger sequencing (Source Bioscience). pTwist-AMP High Copy plasmid was purchased from Twist Bioscience and was used as a vector for all overexpression via transient transfection. Sequences of all LUC7L3 constructs can be found in Supplemental Table 5.

### Cell culture

HeLa cells were obtained from Cell Services at The Francis Crick Institute and cultured in DMEM+GlutaMAX (Thermo Scientific) with 10% FBS and split every 3-4 days using TrypLE Express (Thermo Scientific). Low-passage, wildtype mouse embryonic stem cells (mESCs, 129S8/B6 background) were obtained from the Genetic Modification Service at The Francis Crick institute and cultured feeder free, on Nunc cell culture dishes (Thermo Scientific) coated with 0.1% Gelatin (ES-006-B, Millipore). mESCs were maintained in 2i media^62^, fed every day and split every 2-3 days using Accutase (A6964, Sigma).

All cell transfections were performed using Lipofectamine 3000 (Thermo Scientific) according to manufacturer instructions. Doxycycline induction was performed by changing cell culture media to fresh, pre-warmed media containing 100 ng/mL doxycycline. CLK-IN-T3 (Sigma, SML2649) was dissolved to 5 mM in DMSO. In experiments using CLK-IN-T3 treatments, control cells were treated with the equivalent volume of DMSO.

### Cell line generation

PiggyBac cell lines were generated by co-transfecting HeLa cells with equal amounts of the ePB-ExportReporter plasmid pool and a plasmid expressing the PiggyBac transposase (a gift from Dr. Miha Modic). A transposase-negative control was also prepared. Cells were selected once with 5 µg/mL blasticidin for 4 days, then expanded for one week, before being selected again with blasticidin until all cells in the transposase-negative control were dead. This pool of ePB-ExportReporter positive cells was then expanded and stocks were frozen.

Srrm2 and Son-GFP mESC lines were generated using CRISPR-Cas9 by seeding wildtype mESCs on a 6-well plate and transfecting cells with equal ratios of plasmids encoding Cas9 with puromycin resistance or a donor sequence containing eGFP and approximately 100 bp homology arms complimentary to the C-terminus. Lipofectamine 3000 was used according to manufacturer instructions. The next day, cells were split into 15 cm dishes, allowed to attach, and selected for two days with 1 µg/mL puromycin. Cells were then cultured for 10-14 days until large colonies appeared, before being dissociated and sorted for GFP-positive cells into 96-well plates. After large colonies appeared, clones were split into two 96-well plates. Homozygous clones were screened using PCR of the gDNA from direct lysis of the clones in one plate, and apparent homozygous insertions were confirmed via Sanger sequencing. Successful homozygous clones were expanded and stocks were frozen

### Subcellular fractionation

Nuclear-cytoplasmic fractionation was carried out by trypsinising whole cells and washing twice with PBS. Cells were then pelleted and resuspended in 200 µL cytoplasmic lysis buffer: 50 mM HEPES ph 7.5, 2 mM MgCl_2_, 50 mM 2-mercaptoethanol, 0.025% NP-40, 0.05% saponin, 1X cOmplete EDTA-free protease inhibitors (Sigma). Cells were rotated at 4C for 10 minutes, pelleted, and the supernatant was taken as the cytoplasmic fraction. The nuclear pellet was washed in 1 ml cytoplasmic lysis buffer for 3 minutes, then pelleted again. The supernatant was removed and the nuclei were lysed in 200 µL iCLIP lysis buffer^63^ to obtain nuclear fractions. All centrifugations were performed at 4C and at 300×*g* for 3 minutes. All buffers were kept ice cold. For each fraction, 20 µL was used for western blotting to confirm the efficiency of the fractionation.

### iCLIP

iCLIP was performed according to the iiCLIP protocol^63^, using crosslinked cell lysate containing 1.5 mg of protein per sample. Two replicates were prepared for each CLIP target. For immunoprecipitation, 5 µg of either anti-TRA2B (abcam, ab31353) or anti-GFP (Abcam, ab290) was used. Membrane transfer was performed overnight at 4°C with reduced voltage. For Srrm2-GFP and Son-GFP samples, CLIP membranes were cut into two samples for further processing: low molecular weight (40-300 kDa) and high molecular weight (300+ kDa). For Tra2b samples, the entire lane was cut from 40 kDa upwards.

### iCLIP analysis

CLIP libraries were trimmed and demultiplexed using Ultraplex^64^ and mapped to a small RNA genome containing all rRNA, snRNA, tRNA and snoRNA sequences from GENCODE vM22 using STAR v2.7.0^65^. All unmapped reads were then mapped uniquely to the genome using STAR, with the transcriptome-mapping reads output using the flag --quantMode TranscriptomeSAM and the 5’ end of the read was forced to be aligned using the flag --alignEndsType Extend5pOfRead1. Reads were demultiplexed using UMItools^66^. Crosslink sites were defined as the position immediately upstream of the read.

Metaprofiles around different GeRM regions were created by taking the midpoint of each GeRM region and counting the transcriptome-mapped crosslinks in each position relative to the midpoint. All counts were summed across GeRM regions of the same cluster, then normalised by the number of GeRM regions within the cluster and the number of millions of unique transcriptome-mapping crosslinks in each sample.

### Analysis of public CLIP data

All public CLIP data was processed as described previously^67^, with replicates merged. All crosslinks falling within CDS per sample were counted and the proportion of CDS to total crosslinks was calculated. Samples in the bottom third were excluded. Remaining samples in the bottom third of raw CDS crosslink counts were also excluded. For each possible 5-mer in all CDS sequences, the percentile of the GeRM score was calculated. For each crosslink in each sample, the average percentile of each crosslinked 5-mer was calculated and the mean was taken. If a given 5-mer is crosslinked preferentially in multivalent contexts (relative to the average multivalency of that 5-mer), then the mean GeRM percentile will be high. The mean 5-mer GeRM percentiles for the top 50 most crosslinked 5-mers was compared to all other 5-mers, producing a ratio. This ratio represents the general bias of a protein to bind its preferred motifs in a multivalent context. Only samples with a ratio of at least 1.1 were kept for further analysis.

Only transcripts in the top half of total crosslinks per nucleotide in the CDS across all samples were used for further analysis. For each GeRM region, the ratio of the number of crosslinks falling within that GeRM region compared to the rest of the CDS for that transcript was calculated. The mean of this ratio within each GeRM cluster was calculated for each sample. Samples were then hierarchically clustered and plotted as a heatmap.

### Statistics

All statistics were performed in R 4.2.0. All tests were two-tailed when possible. All pairwise comparisons were made using a Welch t-test unless otherwise stated. When appropriate, p-values were corrected for multiple testing using the Benjamini-Hochberg procedure.

## Supplemental figures

**Figure S1.**
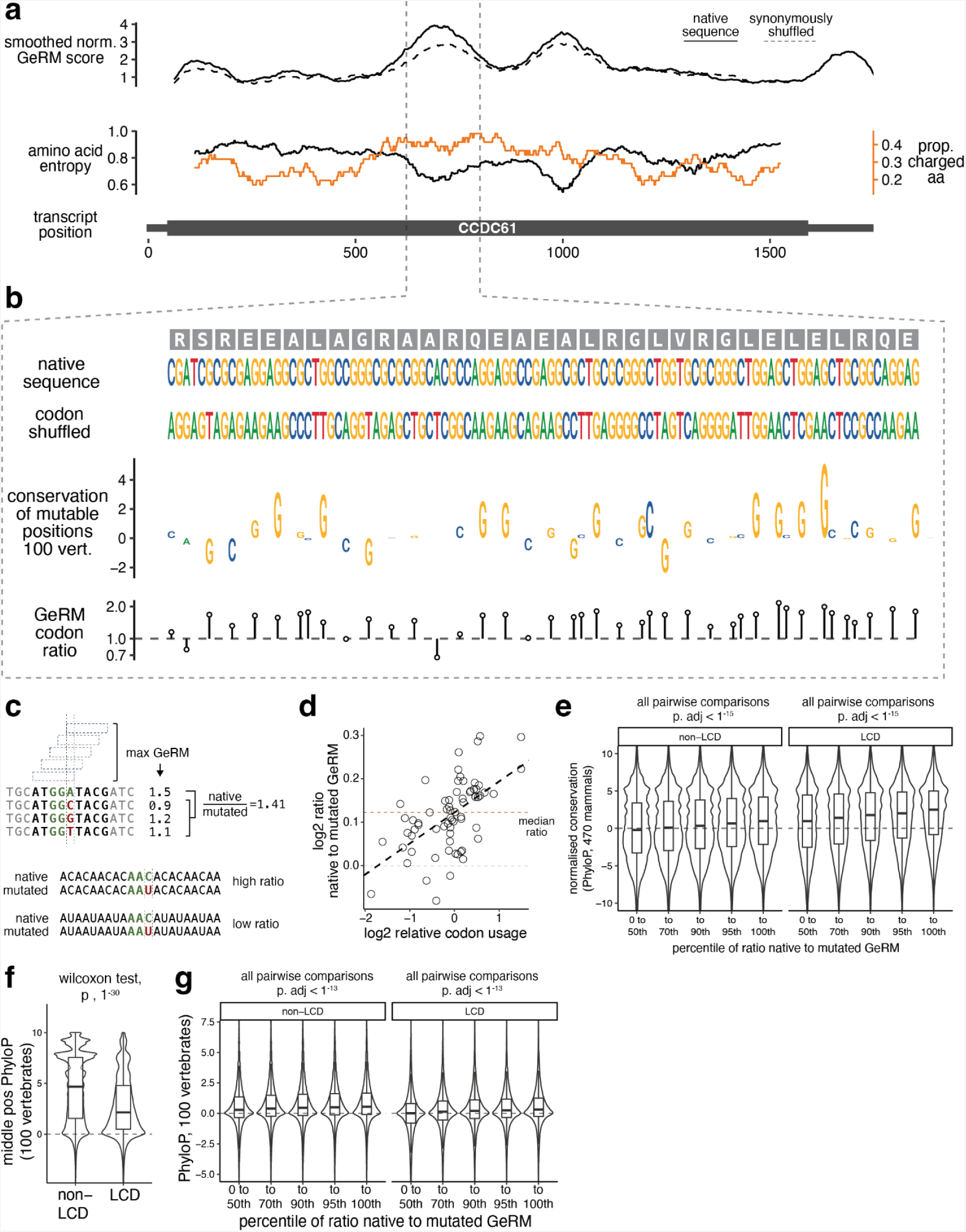
a, An example of a transcript from the gene CCDC61. The smoothed GeRM score is shown along the transcript in the upper panel (solid line), with the dashed line depicting the average smoothed GeRM score after synonymous codon shuffling. The amino acid entropy of the encoded sequence is shown in the lower panel (black line), while the proportion of charged amino acids in that window is shown in the orange line. b, The native DNA and amino acid sequences of CCDC61 within the GeRM peak. A codon shuffled sequence in which all codons must be synonymously shifted is shown. The conservation across 100 vertebrates (PhyloP) for each position that can tolerate synonymous mutation is shown, with the height of the letter corresponding to the PhyloP score. Below, a ratio is shown that compares the GeRM scores associated with the native codon choice to the GeRM scores associated with any possible synonymous mutation. c, a schematic detailing the calculation of the ratio of GeRM between native codon choices and synonymous mutations. d, the mean log2-transformed ratio for each type of possible synonymous mutation. For example, for the arginine codon CGG, position 1 is considered a synonymously mutable position (CGG -> AGG) and position 3 is considered separately as a mutable position (e.g CGG- > CGC). Relative codon usage is calculated by scaling the transcriptomic codon usage such that the mean is 0 and the standard deviation is 1. e, The normalised conservation across 470 mammals of synonymously mutable positions in coding sequences that either encode LCDs or do not. Codons are binned by the degree that the native codon choice supports sequence multivalency, where codons with the highest ratio support the multivalency the most. Conservation values are normalised to the middle position of the codon, which is never synonymously mutable, and each codon is normalised to have a median conservation value of 0. f, PhyloP scores across 100 vertebrates of the middle positions of codons inside or outside of LCDs. g, As in Figures 1H and S1E, but using unnormalised PhyloP scores across 100 vertebrates. Unless otherwise stated, all pairwise statistical comparisons performed using a Welch t-test with FDR correction for multiple testing, where * = p < 1^-15^.

**Figure S2.**
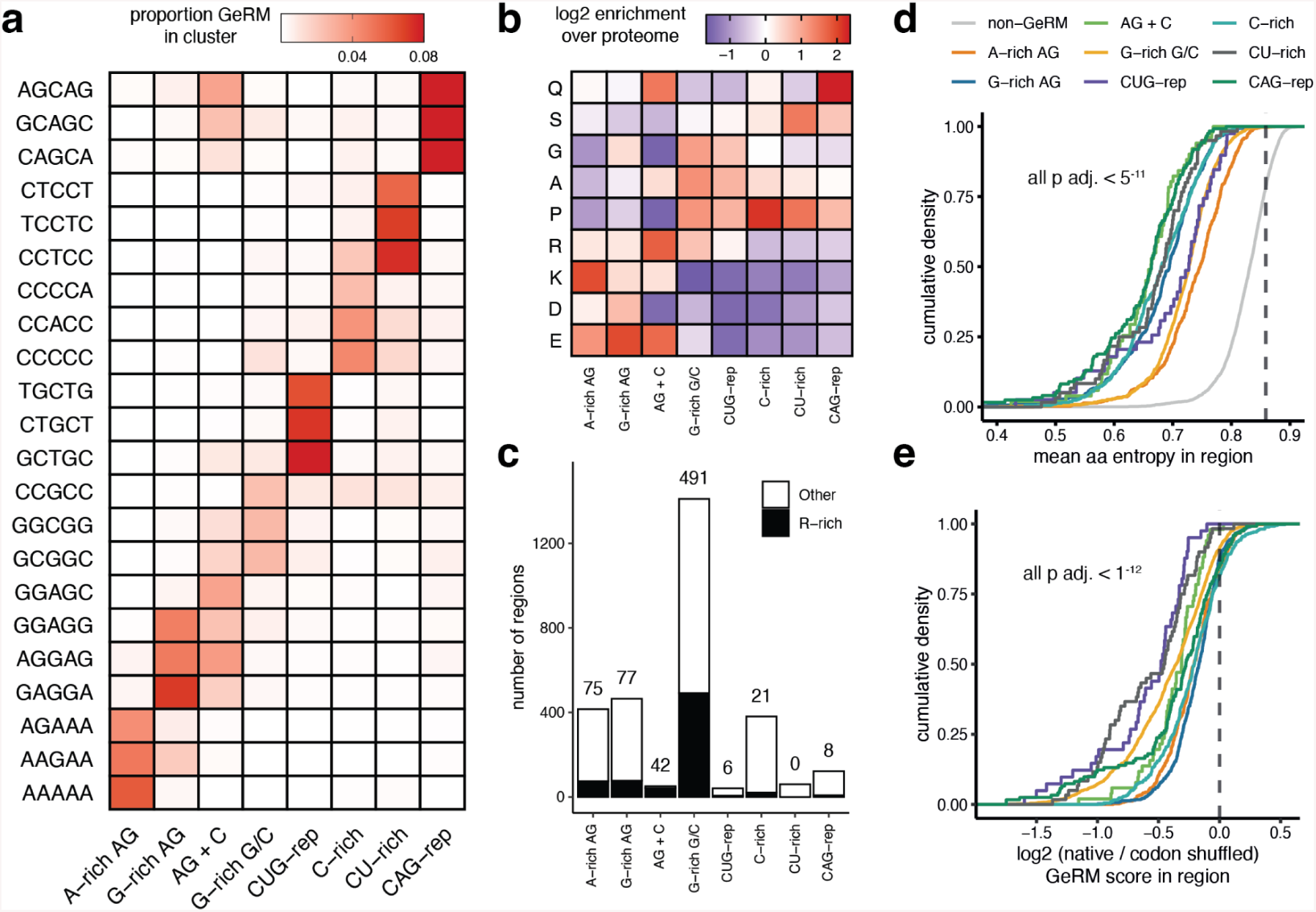
a, the proportion of the total GeRM in each GeRM cluster that is accounted for by a given 5-mer. Only the three most common 5-mers per cluster are shown. b, The relative occurrence of amino acids encoded by different GeRM regions relative to their occurrence in the entire proteome. Only amino acids with notable enrichments are shown. c, The number of GeRM regions per cluster encoding at least two-fold more arginines than expected based on occurrence in the proteome. d, The mean amino acid entropy calculated in sliding windows for amino acids encoded by different GeRM regions or the amino acid sequences encoded by the rest of the CDS (grey line). The dashed line represents the mean amino acid entropy for the entire proteome. Statistical comparisons are Welch t-tests between the GeRM regions and the non-GeRM regions from the same set of proteins, with FDR correction. e, The log2-transformed ratio between the total GeRM calculated from the native sequence within a GeRM region and the mean total GeRM score in the same region when codons have been synonymously shuffled five times within each transcript. Statistical tests are one sample t-tests, with FDR correction.

**Figure S3.**
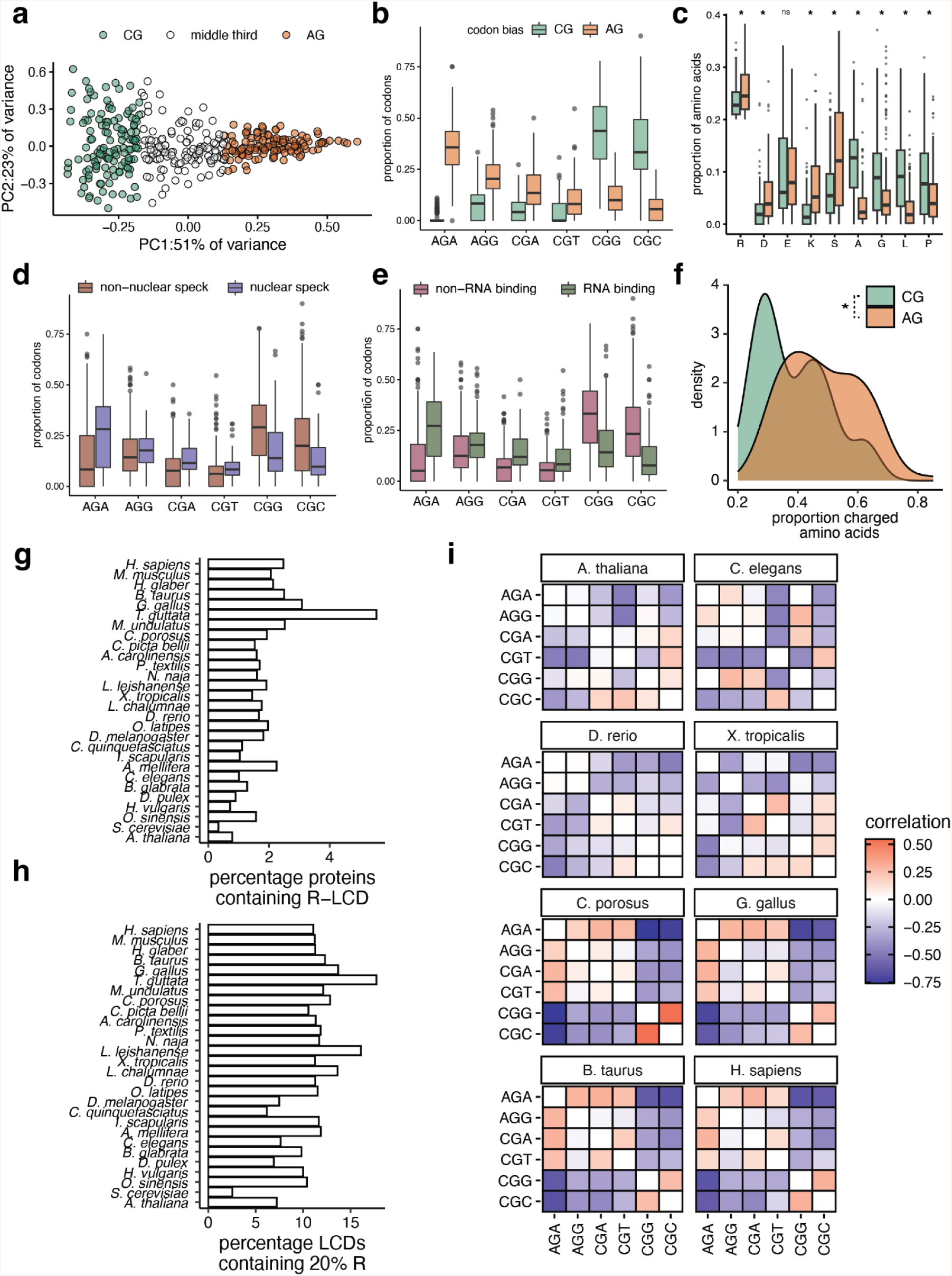
a, Principal component analysis of the arginine codon usage within LCDs containing at least 20% arginine (R-LCDs). R-LCDs are categorised into three groups of equal size using the first principal component. b, The codon usage of the groups of R-LCDs defined in panel a. c, The representation of amino acids within the groups of R-LCDs defined in panel a. d-e, The codon usage of R-LCDs from genes with either ‘nuclear speck’ or ‘RNA binding’ gene ontology terms compared to R-LCDs without those annotations. f, The proportion of charged amino acids within the R-LCD categories defined in panel a. g, The percentage of proteins containing at least one R-LCD in different species. h, the percentage of LCDs that contain at least 20% arginine in different species. i, the Spearman’s rank correlations of codon usage within R-LCDs in different species. All statistical comparisons are Welch t-tests with FDR corrections where appropriate.

**Figure S4.**
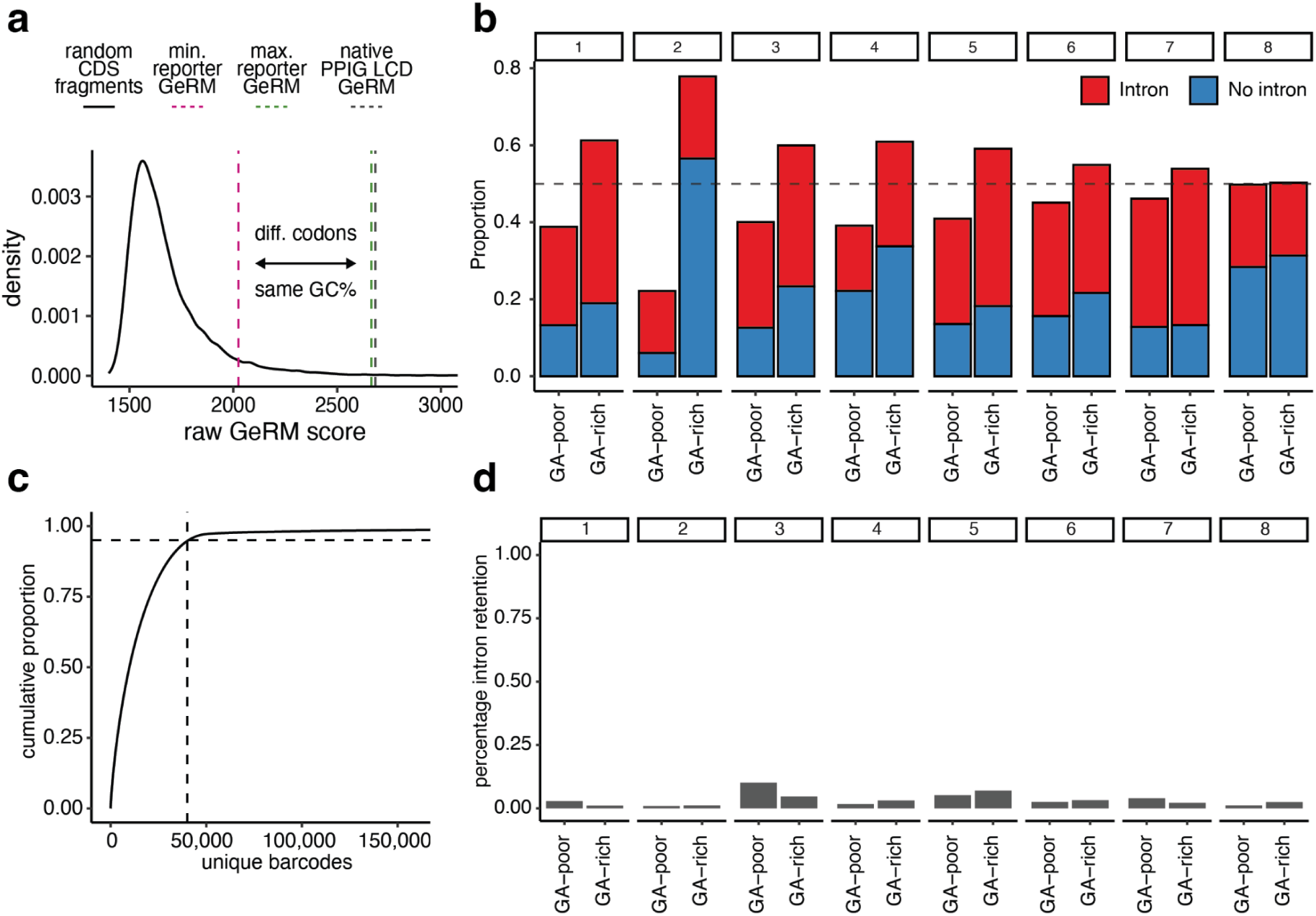
a, The distribution of total GeRM scores within randomly sampled CDS regions of the same size as PPIG_LCD_. The black dashed line shows the GeRM total score of the native PPIG_LCD_ sequence, while the pink and green dashed lines show the minimum and maximum total GeRM scores of the least and most multivalent PPIG_LCD_ CDS from the reporter pool. b, The distribution of different sequence segment types in the reporter plasmid pool as determined by long-read sequencing. c, The cumulative proportion of the total number of reporter barcode reads accounted for by unique reporter barcode sequences in a targeted sequencing experiment. The dashed lines represent the point at which 95% of barcode reads are accounted for. d, The rates of intron retention in reporter transcripts with different CDS codon biases at each position determined from long read targeted sequencing of RNA from cells transfected with the reporter pool for 16 hours.

**Figure S5.**
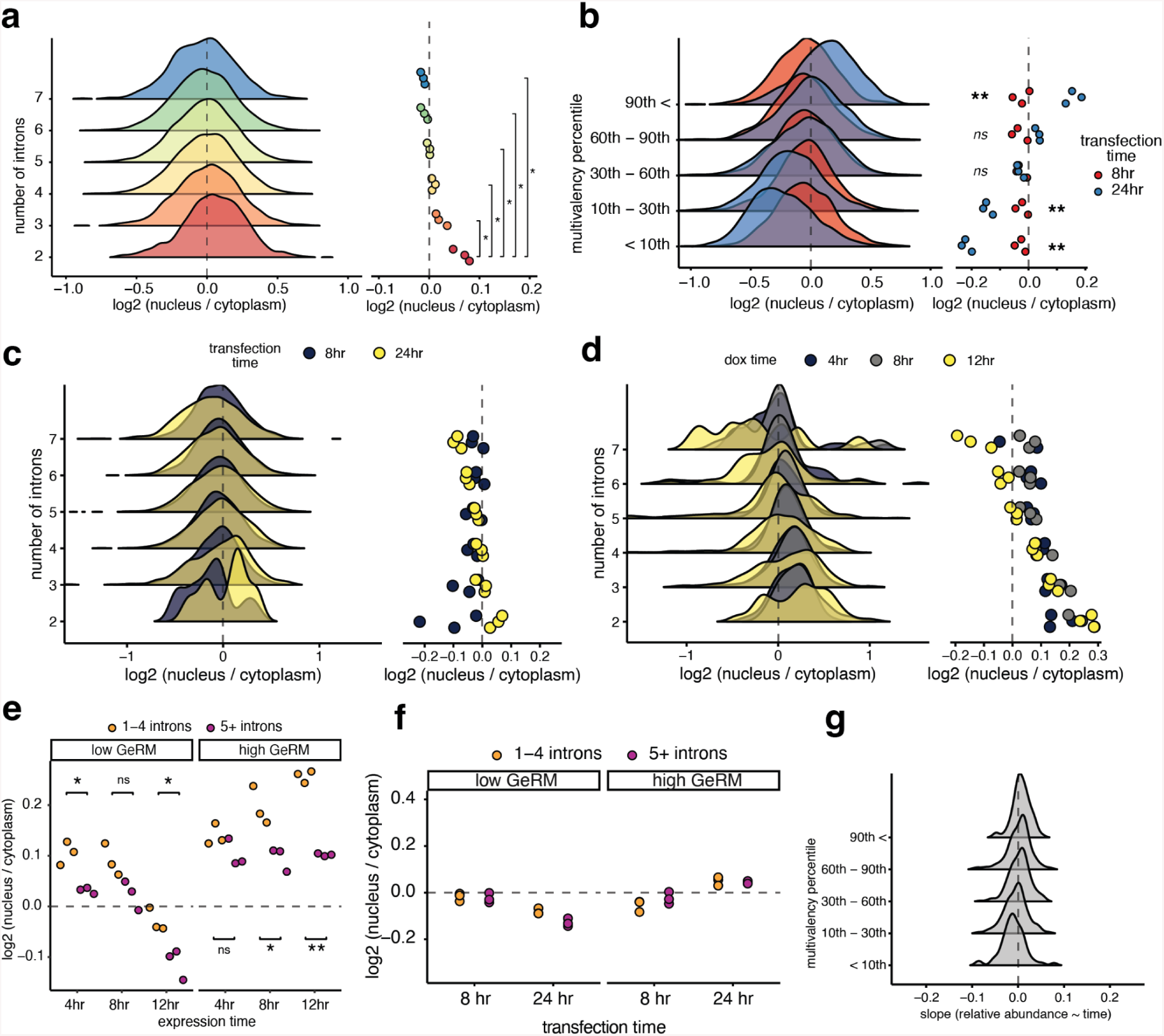
a, The nuclear-cytoplasmic distribution of reporter transcripts binned by the number of introns in their reporter gene. Data from cells transfected with the reporter plasmid pool for 16 hours. b, Nuclear-cytoplasmic distributions of reporter transcripts, but comparing cells transfected for either 8 (short) or 24 hours (long) with the plasmid pool. c, The nuclear-cytoplasmic distribution of reporter transcripts binned by the number of introns in their reporter gene. Data from cells transfected with the reporter plasmid pool for either 8 or 24 hours. d, As in panel e, but from cells in which the reporter library is stably integrated and expressed for either 4, 8 or 12 hours by the addition of doxycycline. e, Nuclear cytoplasmic distributions of transcripts with either above or below average GeRM scores and either 1-4 or 5+ introns in their genes from cells expressing the reporter plasmid pool for either 4, 8 or 12. f, As in e, but from cells transfected with the reporter plasmid pool for either 8 or 24 hours. g, The slopes from regression models of relative reporter transcript abundance over time were calculated by expressing the reporter pool for 16 hours, then collecting fractions at 0, 2, 4 and 8 hours after withdrawing dox. A positive slope indicates that the transcript is relatively more stable in the pool, while a negative slope indicates that the transcript is relatively less stable. All statistical comparisons performed using a Welch t-test with FDR correction for multiple testing, where * = p < 0.05, ** = P < 0.01, *** = P < 0.001.

**Figure S6.**
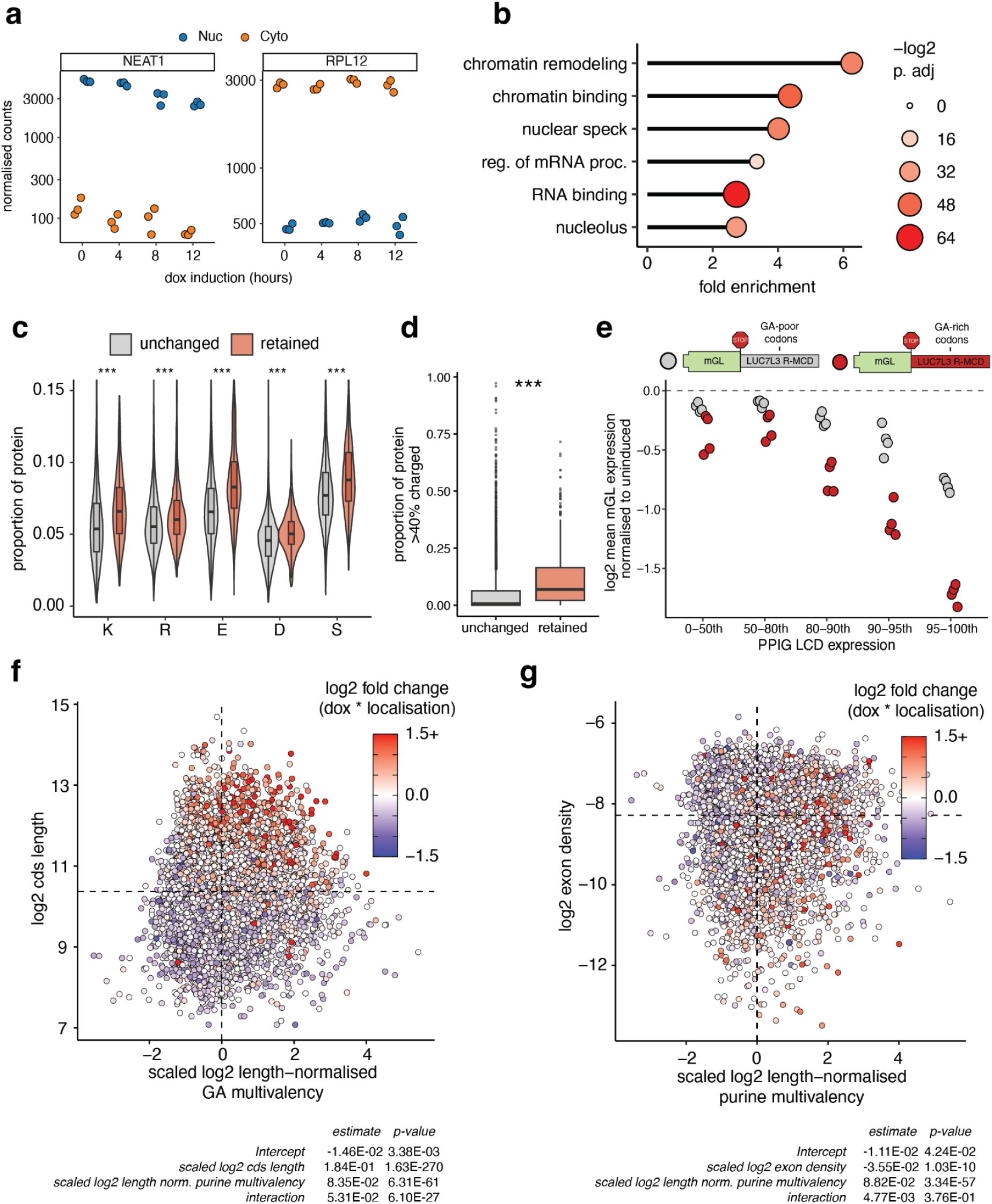
a, Normalised counts for the nuclear lncRNA NEAT1 and the efficiently exported mRNA RPL12 in nuclear and cytoplasmic fractions at different PPIG_LCD_ expression timepoints. b, Gene ontology terms enriched for genes that show a significant nuclear enrichment after 12 hours of PPIG_LCD_ expression. c, The representation of amino acids in proteins encoded by mRNAs that are significantly more retained in the nucleus (with a log-fold change of at least 1) in response to 12 hours of PPIG_LCD_ overexpression compared to mRNAs with unchanged nuclear-cytoplasmic distribution. Statistical comparisons were performed using a Welch t-test with FDR correction for multiple testing, where *** = P < 0.001. d, The same categorisation as in panel c, but comparing the proportion of each protein with at least 40% charged residues calculated in a sliding window of 40 amino acids. e, Flow cytometry measurements of mGreenLantern signal from cells transfected with constructs encoding mGreenLantern with a 3’UTR that is either GA-rich or less GA-rich and with PPIG_LCD_ expressed for 16 hours. Cells are binned based on the percentile of their PPIG_LCD_ expression. mGreenLantern intensities are normalised to samples in which mGreenLantern is transfected but PPIG_LCD_ is not expressed. f, A scatterplot depicting the relationship between CDS length, total GA multivalency (normalised by CDS length) and the shift of an mRNA’s nuclear-cytoplasmic distribution after 12 hours of PPIG_LCD_ expression, where an mRNA that becomes more nuclear after 12 hours is red, and one that becomes more cytoplasmic is blue. The details of the linear regression model, in which CDS length and purine multivalency are used as predictors for the change in nuclear-cytoplasmic localisation, are found below the plot. g, As in panel f, but replacing CDS length with the number of exons per nucleotide length of each transcript.

**Figure S7.**
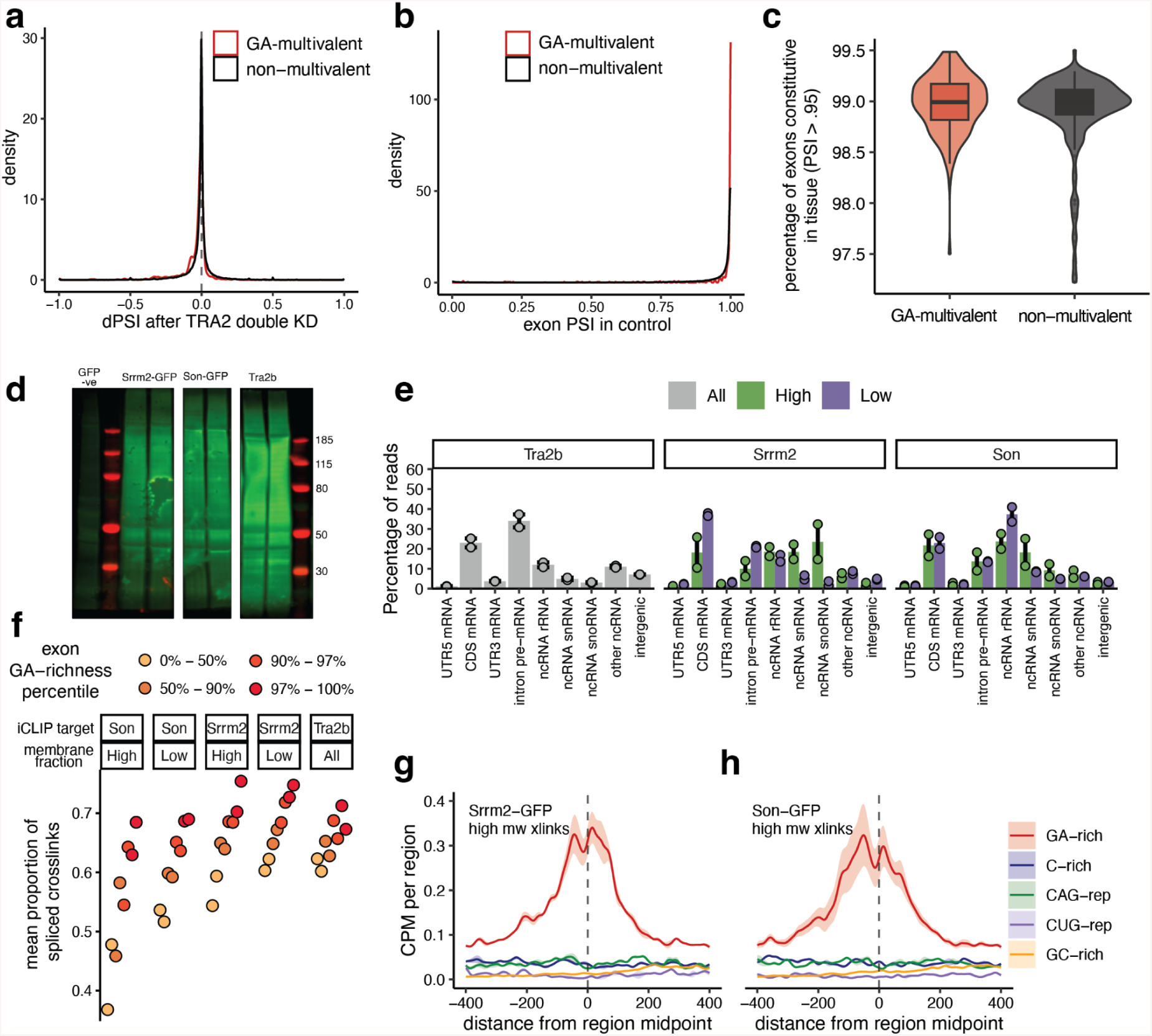
a, Changes in exon proportion spliced in (dPSI) after double knockdown of TRA2A and TRA2B from publicly available data ^29^. Exons are all coding exons from the longest coding transcript per gene with a GA-multivalent region, categorised based on whether the exon overlaps a GA-multivalent region. b, The proportion of exons spliced in (PSI) in the control samples, using the same data and categories as in panel a. c, Using all available human tissues from VastDB, the percentage of exons in each tissue that are constitutive (PSI > .95, exons as defined in panel a). d, An iCLIP membrane showing signal from Srrm2-GFP, Son-GFP and Tra2b iCLIP experiments. For Srrm2-GFP and Son-GFP iCLIP, the membrane was cut in two pieces per sample, separated at approximately 300 kDa. e, The proportion of mappable iCLIP reads mapping to different RNA sub-species. Error bars represent standard error. f, Taking a window of 50 nt from the 3’ end of every internal exon of the longest coding transcript per gene, the proportion of iCLIP reads that contain a splice junction. Exons are binned based on the percentile of their density of GA-containing pentamers. g-h, Metaprofiles of CLIP crosslinks for Srrm2-GFP (high molecular weight fraction) and Son-GFP (high molecular weight fraction) around the central points of subtypes of GeRM CDS clusters in mouse. Solid lines represent the mean across two samples, while shaded regions represent the standard error.

**Figure S8.**
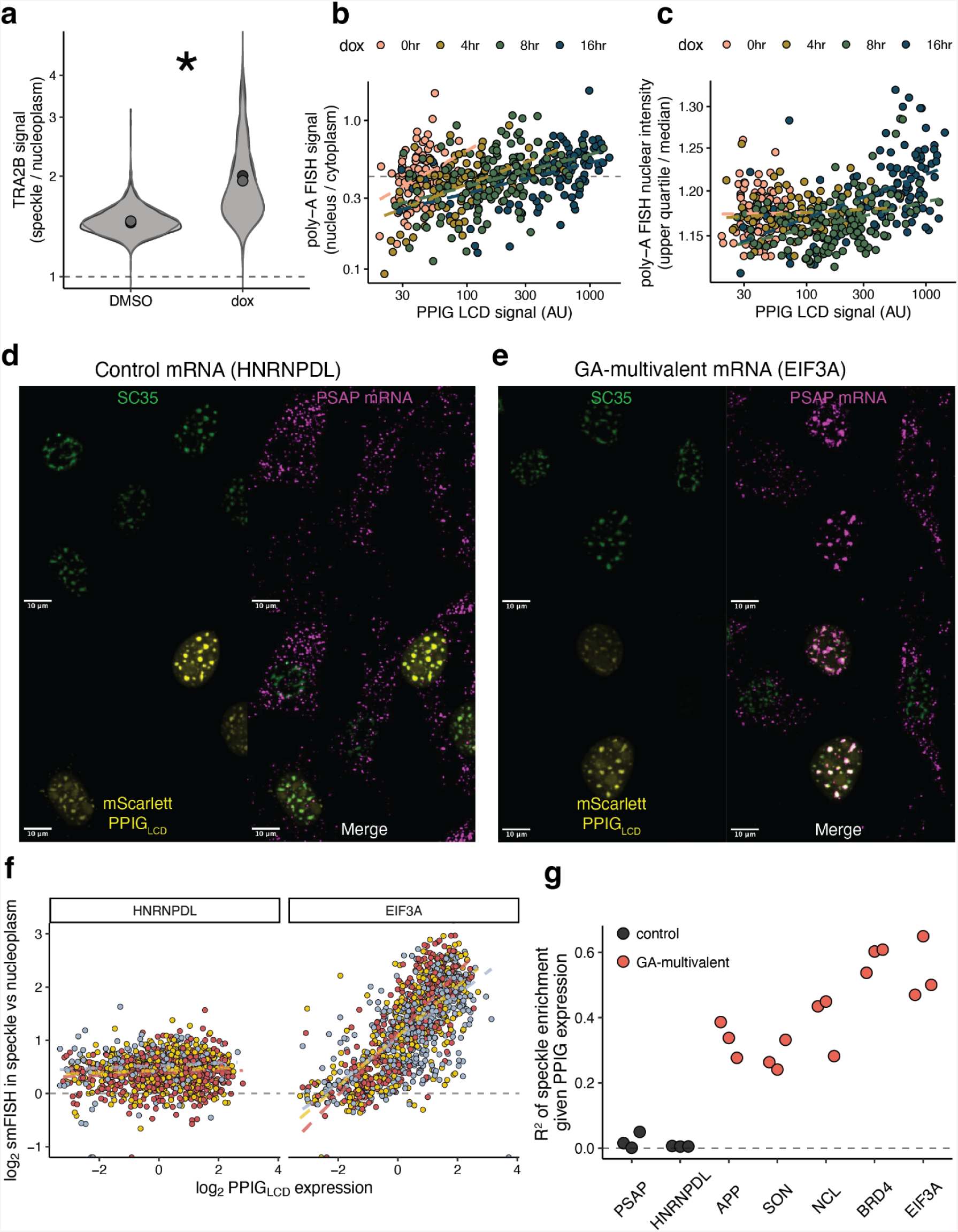
a, The ratio of TRA2B immunofluorescence signal within nuclear speckles or the nucleoplasm in cells treated with doxycycline to induce expression of PPIG_LCD_ or treated with water. The distribution of ratios per nucleus is shown by violin plot and the mean per replicate is shown by the points. Statistical comparison performed on per-repilcate means using a Welch t-test, where * = p < 0.05. b, The ratio of total oligo-dT FISH signal in the nucleus and cytoplasm per cell compared the the intensity of mScarlet-PPIG_LCD_ in the nucleus. Points are coloured based on the length of expression induction with doxycycline. c, The ratio of the upper quartile to the median of nuclear oligo-dT FISH signal as a proxy for the relative intensity of nuclear speckle signal. Ratios are compared the intensity of mScarlet-PPIG_LCD_ in the nucleus, and points are coloured as in panel b. d-e, Example images showing HCR-FISH signal for HNRNPDL (control) or BRD4 (GA-multivalent) mRNAs, with SC35 immunofluorescence and mScarlett-PPIG_LCD_. f, Quantification of the enrichment of HCR-FSH signal within nuclear speckles versus nucleoplasm per nucleus with respect to PPIG_LCD_ expression. Independent replicates are plotted in different colours and regression lines are plotted in dashed lines. g, The fits of linear regression models (R^2^) of the relationship between mScarlett-PPIG_LCD_ expression and mRNA enrichment within the speckle for control and GA-multivalent mRNAs.

**Figure S9.**
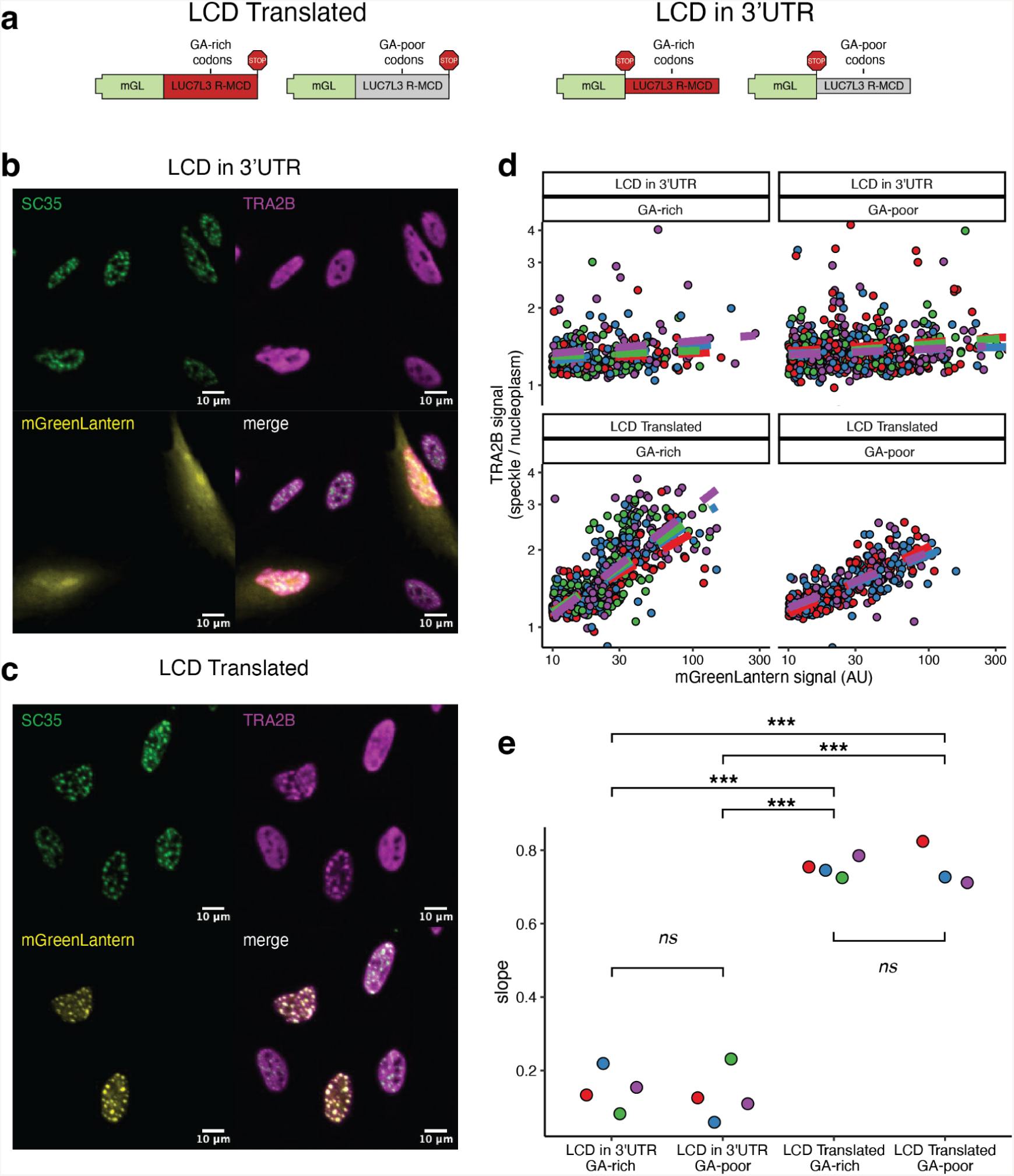
a, A schematic of the mGreenLantern constructs used for the experiments presented in the rest of the panel. b-c, A representative image showing TRA2B and SC35 signal in cells expressing mGreenLantern either with the GA-rich LCD of LUC7L3 as a 3’UTR sequence or as a translated sequence. d, Quantification of the enrichment of TRA2B signal within nuclear speckles versus nucleoplasm per nucleus with respect to mGreenLantern expression for the different constructs shown in panel a. Independent replicates are plotted in different colours and regression lines are plotted in dashed lines. e, The slopes of linear regression models of the relationship between mGreenLantern expression and TRA2B signal within the speckle for the different constructs shown in panel a. All statistical comparisons performed using a Welch t-test with FDR correction for multiple testing, where * = p < 0.05, ** = P < 0.01, *** = P < 0.001.

**Figure S10.**
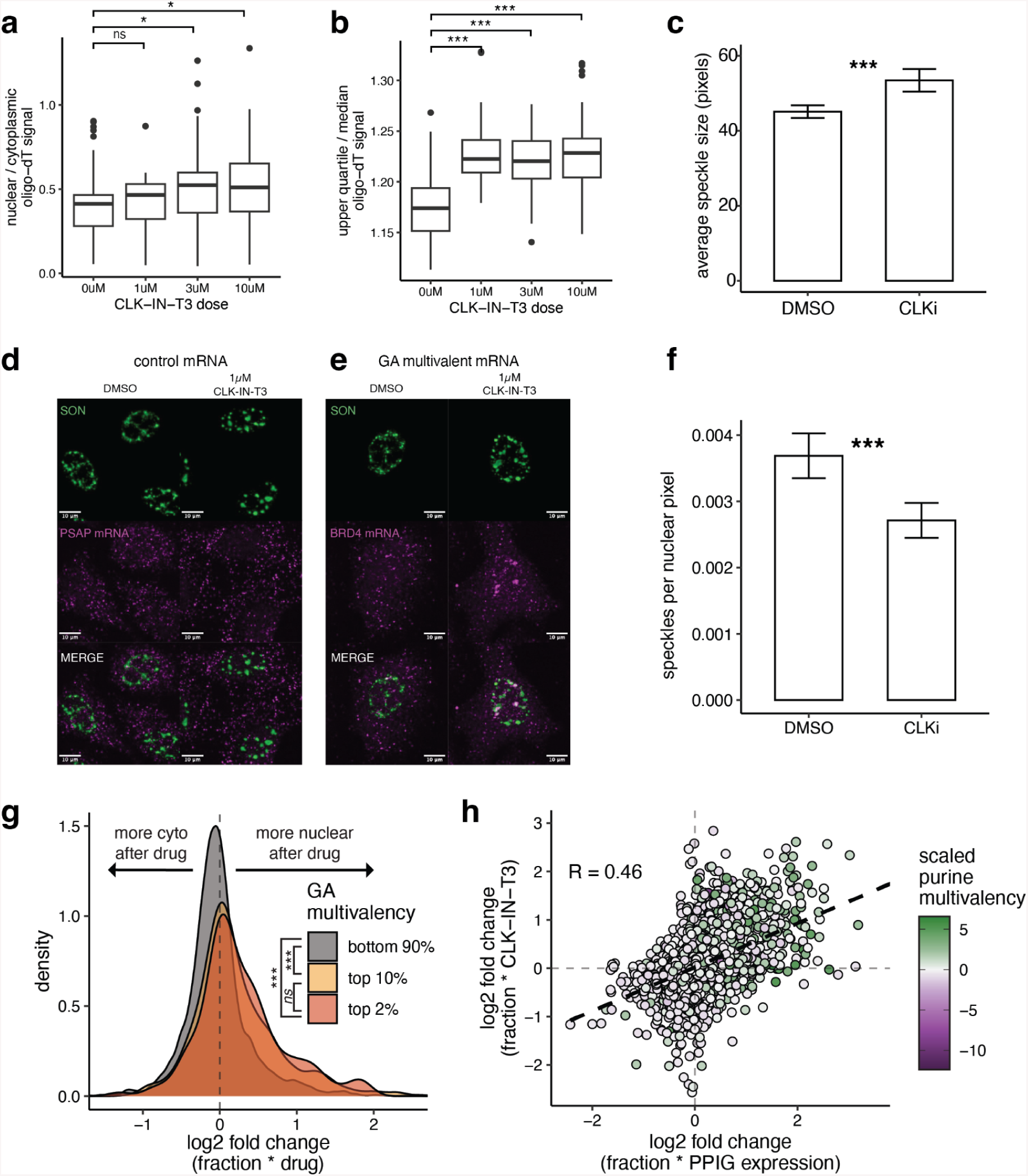
a, The nuclear-cytoplasmic ratio of oligo-dT signal per cell at different doses of CLK-IN-T3 for 8 hours. b, The ratio of upper quartile to median oligo-dT signal per nucleus at different doses of CLK-IN-T3 for 8 hours. c, The average size of speckles per nucleus per replicate in cells treated with either DMSO or 1µM CLK-IN-T3 for 16 hours. d-e, Example image of SON immunofluorescence and HCR-FISH of either control (PSAP) or GA-multivalent (BRD4) mRNA in cells treated either with DMSO or 1uM CLK-IN-T3 for 16 hours. Scale bars are 10µm. f, The number of speckles normalised by nuclear area per nucleus per replicate in cells treated with either DMSO or 1µM CLK-IN-T3 for 16 hours. g, Per gene interaction effect of subcellular fraction and drug treatment from 3’ end sequencing of nuclear and cytoplasmic fractions of cells treated with DMSO or 1uM CLK-IN-T3 for 8 hours. Genes are classified based on the amount of GA-multivalency that they contain in their coding sequences. h, Per-gene log-transformed fold-changes for the interaction terms from PPIG_LCD_ overexpression (12 hours doxycycline) or 1 µM CLK-IN-T3 (8 hours) where a positive fold change means that the gene was more nuclear after treatment. Points are coloured by their scaled purine multivalency. Dashed line denotes the linear model fit to the log-transformed fold changes. All statistical comparisons performed using a Welch t-test with FDR correction for multiple testing, where * = p < 0.05, ** = P < 0.01, *** = P < 0.001.

